# Biomimetic actin cortices shape cell-sized lipid vesicles

**DOI:** 10.1101/2023.01.15.524117

**Authors:** Lucia Baldauf, Felix Frey, Marcos Arribas Perez, Miroslav Mladenov, Michael Way, Timon Idema, Gijsje H. Koenderink

**Author notes:** London Centre for Nanotechnology, University College London, London WCH1 0AW, UK. Institute of Science and Technology Austria, 3400 Klosterneuburg, Austria (Felix Frey). Francis Crick Institute, 1 Midland Road, London, NW1 1AT, UK. Department of Infectious Disease, Imperial College, London W2 1PG, UK. **Author Contributions:** L.B., F.F. T.I. and G.H.K. designed research; L.B., F.F. and M.A.P. performed research; M.M. and M.W. contributed new reagents; L.B., F.F. and M.A.P. analyzed data; L.B. and F.F. wrote the paper; and M.W., T.I. and G.H.K. edited the paper. **Competing Interest Statement:** The authors declare no competing interest.

## Abstract

Animal cells are shaped by a thin layer of actin filaments underneath the plasma membrane known as the actin cortex. This cortex stiffens the cell surface and thus opposes cellular deformation, yet also actively generates membrane protrusions by exerting polymerization forces. It is unclear how the interplay between these two opposing mechanical functions plays out to shape the cell surface. To answer this question, we reconstitute biomimetic actin cortices nucleated by the Arp2/3 complex inside cell-sized lipid vesicles. We show that thin Arp2/3-nucleated actin cortices strongly deform and rigidify the shapes of giant unilamellar vesicles and impart a shape memory on time scales that exceeds the time of actin turnover. In addition, actin cortices can produce finger-like membrane protrusions, showing that Arp2/3-mediated actin polymerization forces alone are sufficient to initiate protrusions in the absence of actin bundling or membrane curving proteins. Combining mathematical modeling and our experimental results reveals that the concentration of actin nucleating proteins, rather than actin polymerization speed, is crucial for protrusion formation. This is because locally concentrated actin polymerization forces can drive a positive feedback loop between recruitment of actin and its nucleators to drive membrane deformation. Our work paints a picture where the actin cortex can either drive or inhibit deformations depending on the local distribution of nucleators.

**Significance Statement:** The cells in our body must actively change shape in order to migrate, grow and divide, but they also need to maintain their shape to withstand external forces during tissue development. Cellular shape control results from an interplay between the plasma membrane and its underlying cortex, a shell composed of crosslinked actin filaments. Using cell-free reconstitution and mathematical modelling, we show that minimal biomimetic actin cortices can mechanically rigidify lipid vesicles while at the same time driving membrane protrusion formation. Our observations suggest that the spatial distribution of actin nucleation determines whether the actin cortex drives or inhibits membrane deformations.

## Introduction

The actin cortex is a dynamic and tightly regulated cytoskeletal structure that supports the plasma membrane of animal cells (1). It combines two seemingly opposing functions, acting as a rigid scaffold that confers mechanical stability to the cell surface, while also actively generating forces that allow the cell to change its overall shape or form local protrusions. The cortex is a thin network of interpenetrating actin filaments that have different structural and dynamic properties. Linear and relatively long filaments (∼ 1 µm) are nucleated mostly by formins (2, 3), whereas branched networks of shorter filaments (∼ 100 nm) are nucleated by the Arp2/3 complex (3) upon activation by nucleation promoting factors such as N-WASP or WAVE (4, 5). The cortical actin layer determines cell surface mechanics depending on the architecture of the filament network (6). At the same time, the cortex actively turns over within tens of seconds (7). It remains elusive how the stiffening and force generating functions of the actin cortex play out to shape the cell surface.

The complex molecular composition of the actin cortex, with over 200 structural and regulatory proteins, has made it difficult to identify the biophysical principles that govern actin-mediated membrane shaping (8, 9). Efforts have thus been made to reconstitute simplified cell-free systems, where actin networks are polymerized on supported lipid bilayers or on the inner or outer surface of giant unilamellar vesicles (GUVs). Using these systems, it has been demonstrated that single actin filaments cannot withstand large compressive loads (10), but stiffer filament bundles such as those found in filopodia or stereocilia (11, 12) are able to push out membrane protrusions (13, 14).

In cells, however, correlative light and electron microscopy has shown that actin filament bundling proteins are recruited to protrusions only after precursors of the protrusion have already been established (15). Bundling proteins and other accessory proteins such as VASP, formins and myosin X later mediate elongation (15–17). Thus Arp2/3 mediated actin polymerization forces alone may be sufficient to initiate cellular protrusions. One reconstitution study confirmed this, but in the context of thick actin cortices grown on the outside of lipid vesicles, creating a geometry inverse to that of the actin cortex in cells (18). It remains an open question whether polymerization forces generated by an actin cortex with more physiological geometry and thickness can also drive the formation of membrane protrusions. Furthermore, it is so far unclear how the rigidity of actin structures impacts membrane shapes. It is well established that the architecture and turnover dynamics of the actin cortex both strongly modulate cortical mechanics and thus the shape of living cells (7, 19). Cell-free reconstitution has, however, focused mostly on systems without significant actin turnover (13, 20–22), or with little excess membrane area (23, 24), where any membrane re-shaping was difficult to observe.

Here we reconstitute biomimetic actin cortices inside GUVs to better understand how rigid, yet dynamic actin cortices may shape the vesicle membrane. We show that branched cortices nucleated by the Arp2/3 complex can stabilize out-of-equilibrium GUV shapes over long timescales of many minutes. At the same time, the actin cortices were also able to drive the formation of membrane protrusions, which often occur in large bouquets, likely as a result of membrane pinning in the dense actin cortex. We show that membrane protrusion formation is strongly dependent on the surface density of the nucleation promoting factor VCA. Protrusion formation is favoured most at intermediate VCA concentrations, which we rationalize by a mathematical model that explicitly accounts for the autocatalytic actin nucleation mechanism of Arp2/3.

## Results

### Membrane-nucleated actin networks mimic cellular actin cortices

To study how actin cortices shape lipid bilayer membranes, we reconstituted a simplified biomimetic actin cortex on the inner surface of GUVs using an emulsion transfer method (25). To selectively restrict actin nucleation to the membrane as in cells (26), we used a 10xHis-tagged VCA construct that binds to nickel-chelating lipids in the membrane. VCA activates the Arp2/3 complex, which in turn allows Arp2/3 to nucleate branched actin cortices (Fig. 1A). The method produces a high yield of GUVs with a broad size distribution centered reproducibly around 9 µm and extending up to 40 µm (25), which covers the lower end of the 10-100 μm size range of mammalian cells (27). Actin was strongly localized to the membrane in almost 90% of the GUVs (Fig. 1B). Some GUVs, however, had a cytosolic actin signal (Fig. 1B, yellow arrow) or lacked any actin signal (Fig. 1B, white arrow), likely because of some variability in the encapsulation efficiency (25). In the following, all analysis refers to GUVs in which we observed membrane-localized actin signal.

**Figure 1.**
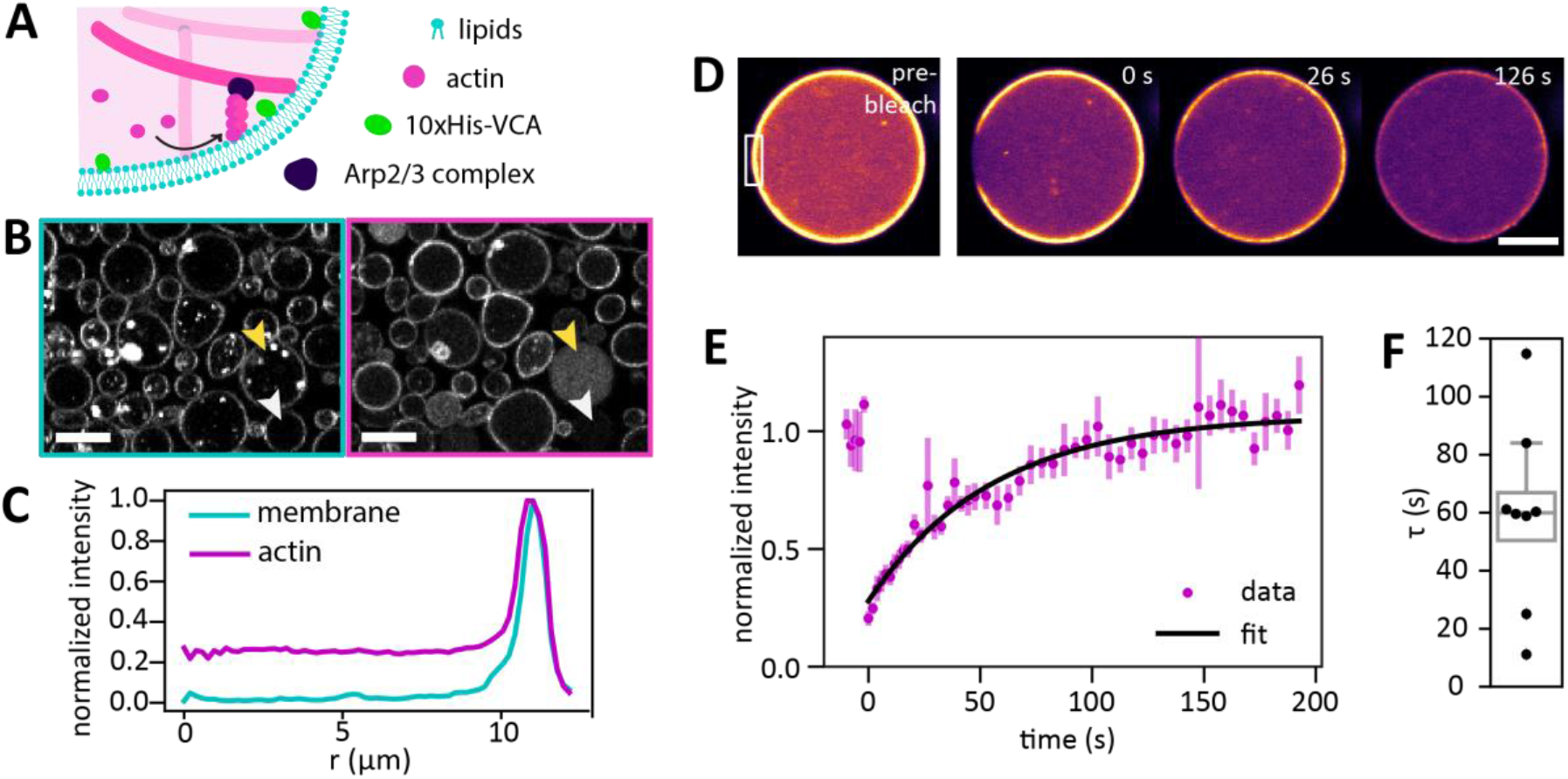
Formation of biomimetic actin cortices in Giant Unilamellar Vesicles (GUVs). (A) Branched actin cortices are nucleated at the inner leaflet of the lipid bilayer membrane by recruiting 10xHis-VCA to the membrane with Ni-NTA-lipids, where it activates the Arp2/3 complex. Activated Arp2/3 nucleates daughter filaments from pre-existing mother filaments that form spontaneously. (B) Confocal images show that actin (magenta) co-localizes in a thin layer with the membrane (cyan) in almost all vesicles. Some GUVs are empty (white arrow) or contain actin in the lumen (yellow arrow). The membrane channel reveals the presence of some bright lipid aggregates. (C) Radially averaged line profiles of the fluorescence intensity of actin and the membrane in the equatorial plane of a GUV show that the majority of the actin signal is concentrated at the periphery within a thin layer whose width cannot be optically resolved. (D) Confocal images taken during FRAP measurements of the turnover time of the actin cortex. Actin is shown in false color (magma) for clarity, with brighter colors indicating higher actin signal. The white rectangle in the pre-bleach image marks the region that was photobleached at *t* = -2 s. Note that the entire vesicle becomes darker over time due to photobleaching during imaging. (E) FRAP curve of the same GUV, showing the fluorescence intensity in the FRAP region corrected for photobleaching during acquisition. Error bars denote the standard deviation, the solid black line shows an exponential fit with a timescale *τ* = 60 s. (F) Characteristic recovery timescales in 9 GUVs from 2 separate experiments, with a mean of *τ* = 59 ± 32 s. Scale bars: 10 μm.

To assess whether the thickness of the reconstituted cortices matches the 150-400 nm reported for the actin cortex of living cells (28–30), we computed the radially averaged intensity profile from confocal images of the actin fluorescent signal in the equatorial plane of the GUVs. Intensity profiles confirmed that most actin signal was indeed concentrated in the periphery of the GUV (Fig. 1C). A lower, homogeneous actin signal was also observed in the GUV lumen, most likely originating from a small cytosolic pool of actin and incompletely removed free dye molecules as verified by fluorescence correlation spectroscopy (Fig. S1). The peak in actin intensity was equally narrow as the peak in membrane intensity, indicating that the thickness of the actin cortex is smaller than the optical resolution of the confocal microscope (< 300 nm). This is in line with a simple estimate of the maximum actin filament lengths (∼300 nm) based on the assumptions that each Arp2/3 complex nucleates one actin filament and all proteins are encapsulated stoichiometrically.

A key characteristic of the cellular actin cortex is the ATP-driven turnover of actin filaments (7). Reconstitution of Arp2/3-nucleated actin cortices has been reported in a few previous studies, but turnover was either not characterized (14, 24, 31) or only assessed qualitatively (23). We therefore measured the dynamics of our cortices by fluorescence recovery after photobleaching (FRAP), similar to approaches used to evaluate actin turnover in living cells (7, 32). We bleached a small rectangular region of the cortex (Fig. 1D, white rectangle) and observed that this region recovered its fluorescence over time (Fig. 1D). Importantly, the width of the bleached region did not change significantly over time (Fig. S2). This indicates that the fluorescence recovery was dominated by exchange of actin monomers or filaments with the GUV lumen via turnover, rather than by lateral diffusion of actin along the membrane. To extract the turnover timescale, we corrected the data for imaging-induced photobleaching and fitted the fluorescence recovery in the bleached region with an exponential recovery with characteristic time *τ* (see Fig. S2 for details on the procedure). This exponential function describes the data well (Fig. 1E), with one dominant recovery timescale of *τ* = 59 ± 32 s (Fig. 1F, *N*=9 GUVs from 2 samples). Strikingly, this timescale is comparable with the ∼20-50 s cortical actin turnover times measured in living cells (7, 33) even though our minimal reconstituted cortices lack proteins that promote actin debranching, filament severing and disassembly (reviewed in (5, 34)). Meanwhile the lipid bilayer membrane remained fluid in the presence of an actin cortex, with a fluorescence recovery timescale of *τ* ≈ 4.9 s (Fig. S3 A, B).

### Branched actin cortices reshape and mechanically stabilize lipid vesicles

Prompted by prior reconstitution studies showing that actin filaments can only reshape lipid membranes at low membrane tension (13, 14, 18), we produced osmotically deflated GUVs. The GUVs had an estimated 7.5% excess membrane area compared to the area needed to cover a sphere of the same volume (35). For deflated vesicles with a fluid surface, we expect smooth and symmetric equilibrium shapes (36). Strikingly, however, most of the cortex-bearing GUVs were frozen in highly irregular shapes, showing sharp corners (pink arrows in Fig. 2A), bleb-like protrusions (yellow arrows), long and thin protrusions (cyan arrows), or highly anisotropic shapes (yellow asterisk). The deformed GUVs were present from the first moment we could image them (∼20 min after GUV formation) and time-lapse imaging revealed that GUV shapes often remained unchanged over many minutes (Fig. S4). Moreover, cortex-bearing GUVs showed practically no thermal membrane fluctuations, in contrast to empty GUVs that fluctuated significantly on the time scale of seconds (Fig. 2B, Movies S1 and S2, additional data in Movie S3). Quantification of the membrane displacements showed that cortex-bearing GUVs were around 3 times less deformable over 50 s than empty GUVs (Fig. 2C).

**Figure 2.**
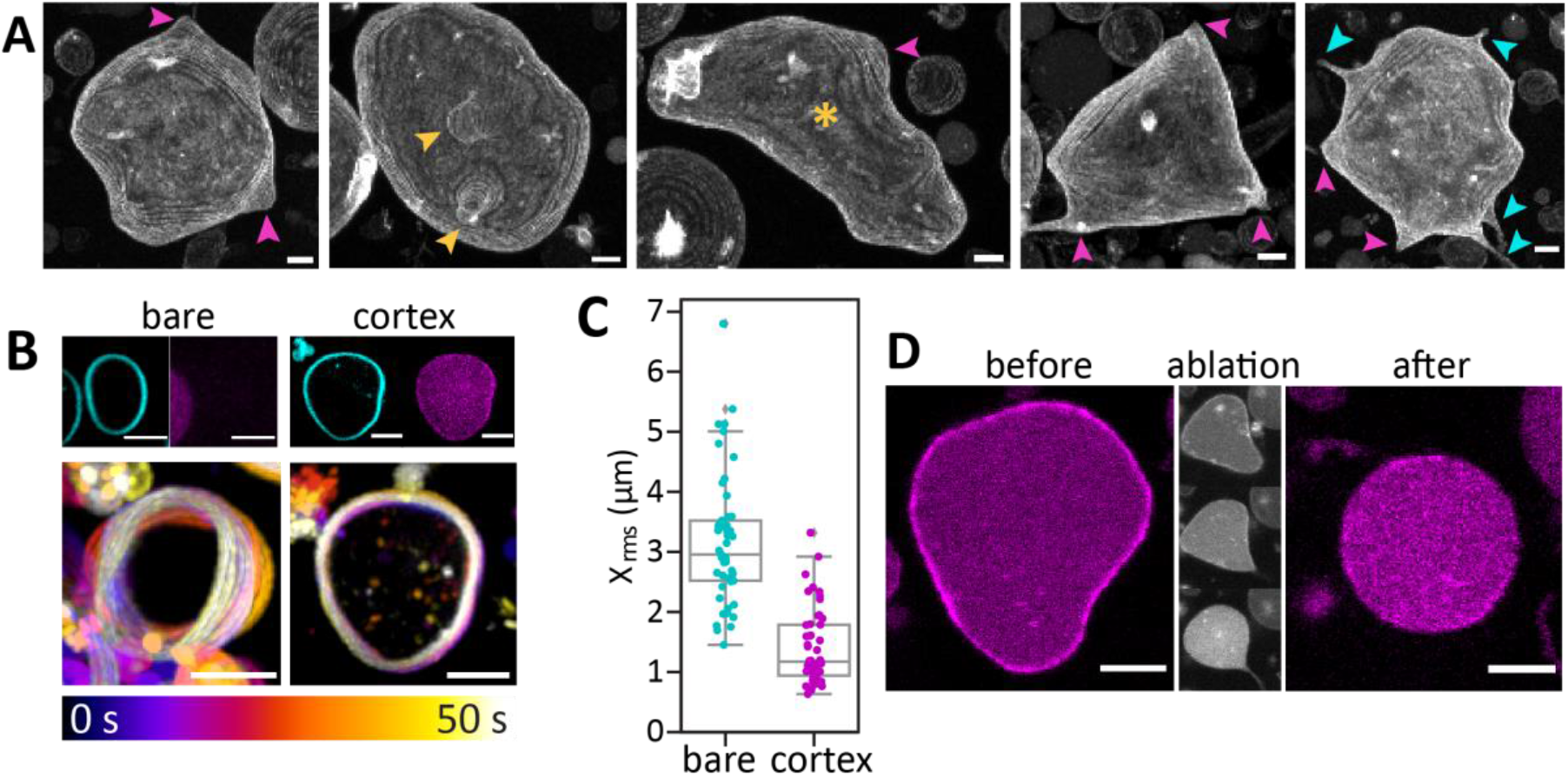
Branched actin cortices stabilize nonequilibrium GUV shapes. (A) Gallery showcasing the diverse GUV shapes in the presence of an actin cortex. Images show maximum intensity projections of confocal z-stacks at 1 μm step height. GUVs exhibit sharp corners (pink arrows), rounded bleb-like protrusions (yellow arrows), long and thin protrusions (cyan arrows) or irregular elongated shapes (yellow asterisk). (B) Actin cortices suppress membrane fluctuations. The bottom row shows color-coded representations of 50 s long time-lapse videos for an empty GUV (left) and a cortex-bearing GUV (right) found in the same sample. The top row shows confocal images with the membrane in cyan and actin in magenta. (C) Root mean square displacement of the membrane between consecutive frames with a 1 s interval for the same GUVs as in B, with a median *X*_rms_ of 3 μm for the empty GUV and 1.2 μm for the cortex-bearing GUV. (D) Photo-ablation of the entire actin cortex returned vesicles to the spherical shapes expected for bare membranes. First and last image: equatorial plane of a GUV before and after ablation for 8 s with a 1.6 W UV laser. Middle column: the same GUV after 1, 4 and 8 s of UV illumination (top to bottom). Scale bars: 5 μm.

To test the role of the actin cortex in membrane shape stabilization, we fragmented the actin cortex by laser ablation, which is widely used to disrupt the actin cortex in cells and has also been used to disrupt actin bundles in GUVs (22, 37, 38). Illumination with an intense UV laser caused GUVs to round up and tended to produce a thin membrane tube from the excess membrane area (Fig. 2D, center-bottom panel). In the example in Fig. 2D, the GUV was spherical after photoablation, but retained a thin, actin-filled membrane tube (right panel). The actin signal in the GUV lumen increased upon ablation, indicating that the cortex was disassembled. Likewise, actin depolymerization with cytochalasin D also returned deformed GUVs to spherical shapes (Fig. S5). Together, these experiments show that branched actin cortices can mechanically stabilize GUV shapes far from the equilibrium shape for a bare lipid bilayer.

### Active actin cortices drive cell-like membrane protrusions

As mentioned above, many cortex-bearing GUVs exhibited long and thin membrane protrusions reminiscent of cellular actin-driven protrusions such as filopodia. We observed two types of outward-pointing membrane protrusions: tubes, whose width was too narrow to optically resolve (Fig. 3A), and spikes, whose width at the base was resolvable (>500 nm, Fig. 3B). Both tubes and spikes were often longer than the GUV diameter but were almost always limited to base widths below 3 µm, regardless of GUV size (Fig. S6 A-C). Protrusions were present in up to 50% of the cortex-bearing GUVs, whereas in control vesicles without an actin cortex, we only observed two tubes and no spikes across 60 GUVs (Fig. 4D, Fig. S7 A). No protrusions occurred when actin polymerized in the GUV lumen, showing that Arp2/3-driven actin polymerization at the membrane drives protrusion formation (Fig. S7 B). Photoablation of spikes using UV-illumination confirmed that actin was needed to maintain protrusions: ablation at the base of spikes either led to complete spike retraction (Fig. 3C, left) or left behind a membrane tube with the site of photoablation collapsed down to a width below optical resolution (Fig. 3C, right, and Fig. S6D). Strikingly, local photoablation of the actin cortex at the base of a spike often also led to shape changes further away, as illustrated in Fig. 3C, where the magenta arrows denote shape changes. Apparently, local cortex ablation can remove shape constraints on the entire cortex.

**Figure 3.**
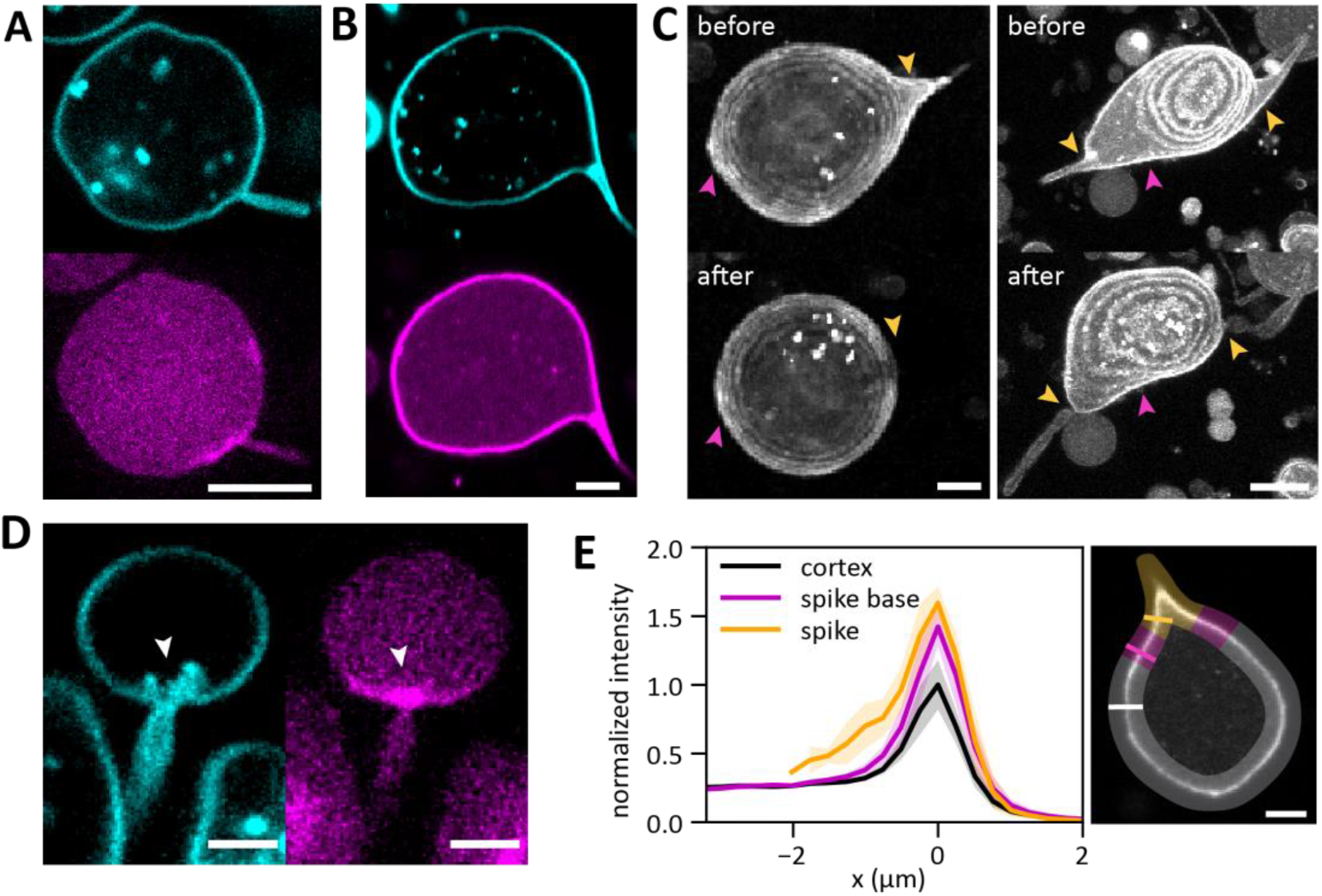
Cortex-bearing GUVs have cell-like membrane protrusions. Confocal images of GUVs with tube-like (A) and spike-like (B) membrane protrusions. Top row shows the membrane (cyan), bottom row the actin (magenta). (C) Maximum intensity projections of two GUVs before (top) and after (bottom) local laser ablation of the actin network in a rectangular region at the spike base (yellow arrows). Destruction of the actin cortex caused retraction of the spike (left) or collapse of the spike to a width below optical resolution (right). Ablation of the spike bases also often caused deformations elsewhere in the GUV (magenta arrows). (D) Confocal image showing the fuzzy membrane signal (cyan, white arrow) associated with a dense patch (magenta) observed at the base of most membrane protrusions. (E) Actin intensity line profiles across the cortex outside the spike (black), at the spike base (magenta) and in the spike (yellow). Lines and shaded regions indicate the means and standard deviations of 6 line profiles for spike and spike base, and 12 line profiles for the cortex on the same GUV. The GUV lumen was located at *x*<0 and membrane at *x*=0, and actin intensities were normalized such that the maximum in the cortex is one. Right panel: Individual intensity profiles were extracted from a confocal image along lines perpendicular to the membrane (solid lines) in the three regions (shaded yellow for the spike, magenta for the spike base, and grey for the rest of the cortex). Scale bars: 5 μm.

**Figure 4.**
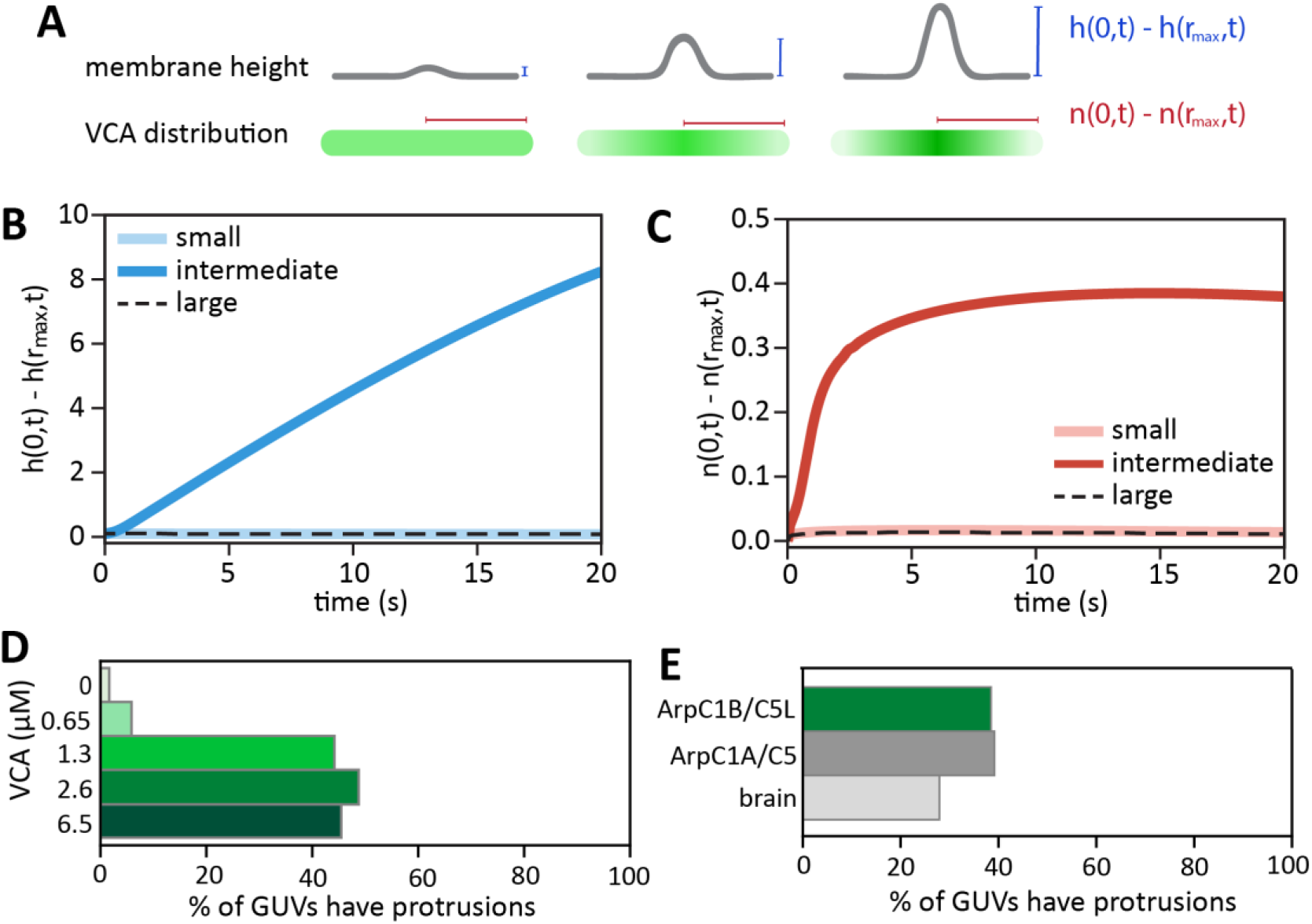
Mathematical modeling shows that the prevalence of protrusions depends on the density of actin nucleators at the membrane. (A) Illustration of the model, where an initial deformation (height *h*) grows over time and changes the initially uniform VCA distribution *n*. (B) Evolution of the height difference between the center of the protrusion (*r=0*) and the boundary of the simulation box (*r=r*_*max*_) for small (*n*_*s*_*=0*.*1*, light blue line), intermediate (*n*_*i*_*=0*.*5*, dark blue line) and large (*n*_*l*_*=0*.*9*, dashed black line) density of VCA on the membrane. (C) Corresponding evolution of the VCA concentration difference between the center of the protrusion and the edge of the bounding box. (D) Bar plot of the fraction of GUVs that were experimentally observed to have tubes or spikes, as a function of VCA concentration. *N =* 60, 85, 129, 121 and 99 GUVs for 0, 0.65, 1.3, 2.6 and 6.5 μM VCA, respectively. (E) Bar plot of the fraction of GUVs with protrusions for three different Arp2/3 isoforms, at a fixed VCA concentration of 2 μM. Data are shown for the human Arp2/3 isoform ArpC1B/C5L (green), which is used in all experiments from Fig. 1-3, for the human Arp2/3 isoform ArpC1A/C5 (dark gray), and for porcine brain Arp2/3 (light grey). *N =* 78, 91 and 111 GUVs, respectively.

To better understand how actin polymerization drives the formation of membrane protrusions, we characterized its spatial distribution in spike-like protrusions. In most cases, actin was visibly enriched in a patch of enhanced actin density at the base of the protrusion (in 78% of 154 GUVs, Fig. 3D). Accompanying actin patches, we often observed an inward-pointing membrane signal whose structure we could not optically resolve (white arrow in Fig. 3D). Line profiles perpendicular to the GUV membrane revealed that the actin cortex was thin and well-defined inside spikes (Fig. 3E). The cortex was slightly thicker and more intense than in the rest of the GUV, but did not fill the spike volume completely.

### Actin polymerization determines the conditions for protrusion initiation

We turned to mathematical modelling to understand how actin polymerization initiates membrane protrusions. Our model describes the early stages of the three-dimensional (3D) shape evolution of an initially planar membrane under the influence of actin polymerization, and was inspired by the work of Gov and Gopinathan (39, 40). We considered the locally flat membrane patch of a GUV, to which the actin nucleation promoting factor VCA is bound. We treated the local VCA density as a proxy for local actin polymerization, which may drive membrane deformations. We assumed that the temporal evolution of the VCA surface density *n(x,y,t)* is determined by diffusion and autocatalytic actin growth (41):

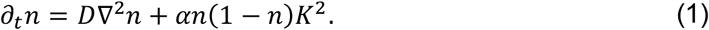

The first term on the right-hand side of Eq. (1) describes the lateral diffusion of VCA on the membrane with diffusion constant *D*. FRAP experiments confirmed that VCA diffused freely on the membrane (Fig. S3 C-D). The second term on the right-hand side of Eq. (1) describes the membrane accumulation of VCA. We assumed that VCA is more likely to reside on the membrane where actin, and therefore VCA, is already present. In addition, we assumed that the accumulation of VCA slows down as the membrane becomes more and more covered with VCA. We therefore chose the term to be proportional to *n(1* − *n)*, so that the VCA number density *n* is normalized and bounded between 0 and 1. Since the VCA density is normalized, the phenomenological ‘enrichment constant’ *α* has the same units as a diffusion constant. Experimentally, we found no high-curvature membrane deformations unless actin was present and polymerizing on the membrane. Thus, wherever the membrane is curved, the probability is high that actin is present at that site. In the model, we therefore posited that the actin (and by extension VCA) density is correlated to the (total) membrane curvature *K*, but independent of whether the membrane is curved inward or outward. The simplest way to accommodate this symmetry is to assume that VCA enrichment is proportional to *K*^*2*^. Note that we did not make any explicit assumptions about curvature generation by actin nucleating proteins, which is often postulated in theoretical works but so far lacks clear experimental evidence (39, 40, 42).

We described the shape of the membrane patch by a ‘height field’ *h(x,y,t)*, which denotes the distance between the membrane position and some flat reference plane. We assumed that the time evolution of the membrane height *h* is determined by the polymerization of actin filaments and the tendency of the membrane to mechanically relax to a flat state,

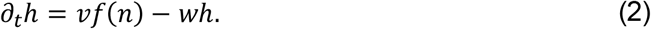

The first term on the right-hand side of Eq. (2) describes how the membrane is pushed outward in the normal direction by polymerizing actin filaments. Here, *v* represents the actin polymerization velocity. The VCA response function *f(n)* describes how the VCA density affects actin polymerization. We chose *f(n)* to have a sigmoidal shape to reflect two reasonable assumptions. First, VCA acts as an actin nucleator, so some minimal concentration of VCA is necessary to induce actin polymerization. Second, actin polymerization saturates at high VCA density as the density of membrane-bound VCA is limited (see ‘Modelling the formation of protrusions’ and Fig. S8, S9, S10 and Table S1 in the Supporting Information). The second term on the right-hand side of Eq. (2) describes how membrane deformations mechanically relax towards a flat state with rate *w*, which is determined by membrane tension and bending rigidity (42). As initial condition, we started from a homogeneous VCA distribution on a membrane patch with a small-amplitude deformation at the origin with a Gaussian shape. Such a deformation may be caused by thermal fluctuations of the GUV membrane (18). Fig. S11 illustrates how the different terms in our model affect the evolution of membrane position and VCA distribution.

We solved the model numerically in polar coordinates, assuming rotational symmetry, for low, intermediate, and high density of VCA on the membrane. Starting from a small initial deformation and uniform VCA distribution, we monitored how the VCA distribution evolved over time, and how the height of the membrane in the center of the membrane patch changed compared to the flat outer boundary of the simulation box (Fig. 4A). For a low initial VCA density, the membrane remained flat (Fig. 4B, light blue line). For a high initial VCA density, the membrane was pushed outward with the same velocity everywhere, thus again remaining flat (Fig. 4B, dashed black line). By contrast, the localized membrane deformation rapidly grew in amplitude for intermediate initial VCA density (Fig. 4 B, dark blue line). Similarly, the difference between the VCA density at the center of the protrusion compared to the outer boundary remained zero for both low and high VCA densities. By contrast, the VCA density in the center of the membrane patch increased relative to the outer edge for intermediate VCA density, reaching a plateau after a few seconds (Fig. 4C). The model thus predicts that the prevalence of thin membrane protrusions should depend non-monotonically on the density of nucleation promoting factors on the membrane.

To experimentally test this prediction, we measured the prevalence of membrane protrusions for GUVs in which we varied the concentration of encapsulated VCA while keeping all other membrane and cortical components the same (Fig. 4D). At low VCA concentrations (1.3 µM, corresponding to an estimated surface coverage of ∼4.5% (see Extended Methods)), we observed very few protrusions. Tubes and spikes became more prevalent at intermediate VCA concentrations, with almost 50% of all GUVs exhibiting thin protrusions. At high VCA concentrations (6.5 µM, corresponding to an estimated surface coverage of 25%), we observed a slight reduction in the prevalence of GUVs with protrusions, with spikes in particular appearing less frequently (Fig. S12). These findings therefore confirm the theoretical prediction that protrusions are promoted by intermediate densities of nucleation-promoting factors.

We also experimentally tested the impact of the nucleation ability of Arp2/3 on protrusion formation, by comparing three different types of Arp2/3 complex: the human Arp2/3 isoforms containing ArpC1B/C5L and ArpC1A/C5 (43) (green and dark gray, Fig. 4E) and the Arp2/3 complex from porcine brain (light gray), which contains a mix of isoforms and is widely used in reconstitution studies (18, 23, 24, 44). Pyrene assays showed that actin filament elongation rates were comparable for the two human Arp2/3 isoforms, with ArpC1B/C5L leading to slightly faster polymerization (82 and 71 nM/s for ArpC1B/C5L and ArpC1A/C5, respectively), but was reduced by a factor of 5 for porcine brain Arp2/3 (15 nM/s) (Fig. S13). The abundance of membrane protrusions was virtually identical in GUVs with the two human Arp2/3 isoforms but was reduced by 30% with the porcine brain Arp2/3 (Fig. 4E). Apparently, protrusion initiation is only weakly dependent on the rate of actin elongation. This observation is consistent with the predictions from our model, where we can vary the actin filament growth rate *v* by 3 orders of magnitude without qualitatively changing the outcome (Fig. S14).

### Membrane protrusions can form in bouquets

In around 10% of cortex-bearing GUVs, we found one or more ‘bouquets’ of membrane protrusions, with multiple protrusions emanating from the same spot (Fig. 5A, B). Bouquets were present irrespective of GUV size (Fig. 5 C), and had up to 11 individual protrusions that fanned out over a wide angular range (Fig. S7 E). The bouquet bases were often associated with a fuzzy membrane signal similar to that of isolated membrane protrusions (Fig. 5C). Also, bouquet bases were almost always strongly enriched in actin (88% of 58 bouquets, Fig. 5D). This observation suggests that bouquets form when actin accumulates at the base of a tube or spike, increasing the local polymerization force density and initiating the formation of a new protrusion. Membrane protrusions in a fluid bilayer membrane are normally expected to fuse into a single protrusion (39). We speculate that membrane pinning in the dense actin network may stabilize the bouquets and prevent fusion. If this idea is true, then VCA should also be enriched in bouquet bases, as it mediates the interaction between the membrane and the actin cortex by binding Arp2/3 and also capping actin barbed ends (45, 46). Indeed, fluorescent labeling of VCA confirmed that this protein was enriched at the base of bouquets (Fig. 5F, G).

**Figure 5.**
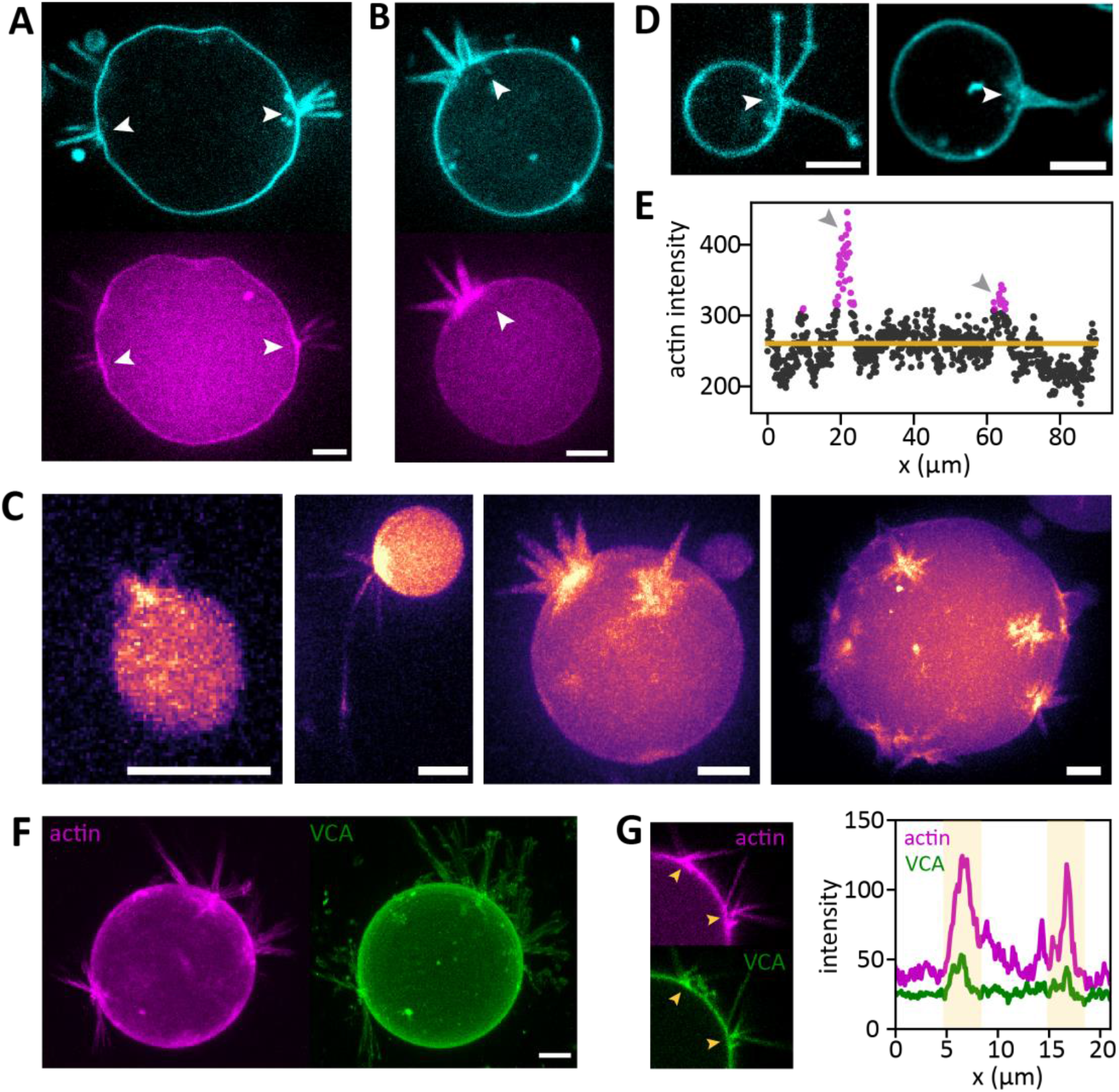
Actin-driven membrane protrusions often occur in bouquets. (A, B) GUVs with bouquets of tubes (A) or spikes (B). Membranes are shown in cyan, actin in magenta, and bouquet bases are indicated with white arrows. The images show maximum intensity projections of three consecutive 1 μm confocal slices. (C) Gallery of different GUVs with bouquets highlights that bouquets occur on GUVs of all sizes (first image: diameter of 5 μm, last image: diameter of 35 μm), with widely varying protrusion lengths, and with either one or multiple bouquets per GUV. Images show maximum intensity projections of z-stacks, smoothed with a 0.5 px Gaussian filter and with actin shown in false color (magma) to better visualize the faint protrusions. (D) Confocal images of the membrane often reveal fuzzy membrane signal (white arrows) at the base of bouquets. (E) Actin intensity profile along the membrane of the GUV shown in (A), clockwise from the top. Gray arrows indicate the positions of the bouquets indicated in (A). Points are highlighted in magenta where the intensity exceeds the mean actin signal (yellow line) by more than 2.5 standard deviations. (F) Confocal images of actin (magenta) and VCA (green) for a GUV with bouquets. Images are maximum intensity projections. (G) Left: Zoom-in on a single confocal image of the GUV shown in F, showing two bouquet bases (yellow arrows). Right: Line intensity profiles along the membrane (clockwise from the top) reveal that VCA and actin are both enriched at the bouquet bases (yellow shaded regions).

## Discussion

To probe how the mechanical interplay between the cell membrane and its underlying actin cortex controls cell shape, we reconstituted minimal branched actin cortices inside cell-sized GUVs. By nucleating actin with Arp2/3 and VCA, we were able to obtain thin and dynamic membrane-anchored cortices with an actin turnover timescale comparable to timescales reported in living cells. The turnover timescale is orders of magnitude faster than the depolymerization of actin filaments in bulk solution, which takes hours (47). The faster turnover inside the GUVs is likely due to the potent nucleation activity of Arp2/3 and VCA (48). At the membrane, G-actin is rapidly incorporated into the cortical network, depleting the GUV lumen of actin monomers and bringing the concentration there below the critical concentration for barbed end elongation. Consequently, any uncapped actin filaments in the GUV lumen will rapidly depolymerize, feeding their constituent monomers back into the cortical layer. The rate limiting step for disassembly of branched actin networks is thought to be debranching (49). Reconstitution studies have shown that debranching takes several minutes in unloaded conditions (50), but piconewton forces will accelerate debranching to well under one minute by causing forced Arp2/3 unbinding from the mother filament (50). The rapid turnover we observe hence suggests that the branches may be under mechanical load. Consistent with this idea, we saw evidence of long-range elastic stress upon local photoablation of the actin cortices. In future studies it will be interesting to study the molecular mechanism of turnover in more detail, for instance by including proteins such as cortactin that modulate branch stability (43, 51, 52).

We found that the actin cortices, despite being thin and dynamic, strongly suppressed thermal fluctuations of the membrane and stabilized nonequilibrium vesicle shapes on timescales up to several hours. While we could not directly observe how the out-of-equilibrium GUV shapes were formed, we speculate that they arise from buckling of the actin cortex due to local inhomogeneities in cortical thickness or mechanical properties. We saw evidence of such inhomogeneities in the form of small actin intensity variations throughout the cortex (Fig. S2 B). Given that actin polymerization is rapid (hundreds of seconds, Fig. S13), direct imaging of the process of actin-mediated vesicle deformation is not straightforward. It will likely require including a mechanism to trigger actin polymerization on demand, for instance by permeabilizing the GUV with pore-forming peptides to allow for content exchange with the outer buffer (23, 53), or by delivering cytosolic contents by targeted membrane fusion (54).

We found that the actin cortices were also able to drive the formation of long and thin membrane protrusions in a manner dependent on the surface density of VCA. The maximum lengths of the protrusions (∼30 μm) far exceeded typical lengths of filopodia, which are usually constrained to lengths of a few micrometers by the diffusion rate of actin monomers and the buckling instability of actin bundles (10, 55). Our system lacks actin crosslinking proteins required for bundling actin filaments, but membrane confinement could potentially give rise to spontaneous actin bundling in the tips of the protrusions (56, 57). Based on the spike retraction observed when we locally ablated actin at the base of the spikes, we speculate that the branched actin cortex keeps the base of the spikes open by exerting outward polymerization pressure normal to the membrane.

To probe the mechanism of protrusion initiation, we compared our experimental observations with the predictions of a mathematical model that explicitly accounts for the autocatalytic actin nucleation mechanism of Arp2/3. Our model predicts that protrusion initiation should be favored at intermediate concentrations of the nucleation promoting factor VCA on the membrane, but suppressed at both low and high VCA concentrations. At low VCA concentration, only single filaments can push on the membrane, which are too weak to initiate a protrusion. At intermediate VCA concentrations, multiple actin filaments can polymerize in close proximity, and due to increased recruitment of VCA to sites where actin is already present, small membrane deformations are stabilized and develop into larger protrusions. At high VCA concentrations, spatial fluctuations in VCA density are suppressed and growth of a continuous network is favored over the formation of local protrusions. Actin barbed-end capping by VCA likely also helps to inhibit protrusion formation at high VCA concentrations (45, 46, 58), although we did not account for this in our model. Consistent with the model’s prediction, membrane protrusions were indeed suppressed at high VCA concentrations. Both the model and experiments with different Arp2/3 isoforms demonstrate that the propensity to form membrane protrusions was only weakly dependent on the actin polymerization rate. We therefore conclude that, while fast actin elongation is beneficial for protrusion formation, the limiting factor is the local VCA availability, suggesting that localized force, not polymerization velocity, is key for generating actin-driven protrusions. Note that VCA not only mediates actin polymerization by activating the Arp2/3 complex, but also mediates actin-membrane adhesion by binding to filament barbed ends (45). To resolve the relative contributions of these two functions, it will be interesting to observe protrusion formation in the presence of the VCA domain of less efficient nucleation promoting factors. Here we used VCA from N-WASP, which has two actin-binding WH2-domains, but VCA domains from other proteins such as WAVE or WASH possess only one WH2 domain and are known to be less efficient in driving actin polymerization through activation of Arp2/3 and in concert with SPIN90 (59, 60).

Our work demonstrates that actin polymerization alone, without any assistance from actin bundling or membrane curving proteins, is sufficient to produce cell-like protrusions from biomimetic cortices. Based on our findings, we can propose a working model for the actin-membrane interactions that shape the GUV surface: First, actin begins to polymerize at the GUV membrane, where it is nucleated by VCA and Arp2/3 (Fig. 6, top). If actin polymerization occurs uniformly across the membrane, then the actin cortex grows homogeneously and stiffens the GUV surface (Fig. 6, left). If, however, a small thermally induced membrane deformation happens to coincide with locally enhanced actin polymerization, more VCA is recruited to the deformation, driving positive feedback that produces a long and thin membrane protrusion (Fig. 6, right). Since large amounts of growing actin filaments accumulate at these protrusion sites, the likelihood increases that they initiate another protrusion in the same area, creating a bouquet (Fig. 6, right bottom). In the context of membrane protrusion formation in cells, our reconstituted system confirms observations in cells suggesting that Arp2/3-driven actin nucleation is sufficient to initiate membrane protrusions (15). The cell-free system we present here provides a powerful platform to dissect the molecular mechanisms of protrusion initiation and elongation. It will for instance be interesting to incorporate temporally controlled recruitment of actin binding proteins to identify how this influences the reorientation of polymerization forces. Further, it will be interesting to compare the mechanical effects of branched cortices studied here with that of cortices made of longer, linear formin-or

**Figure 6.**
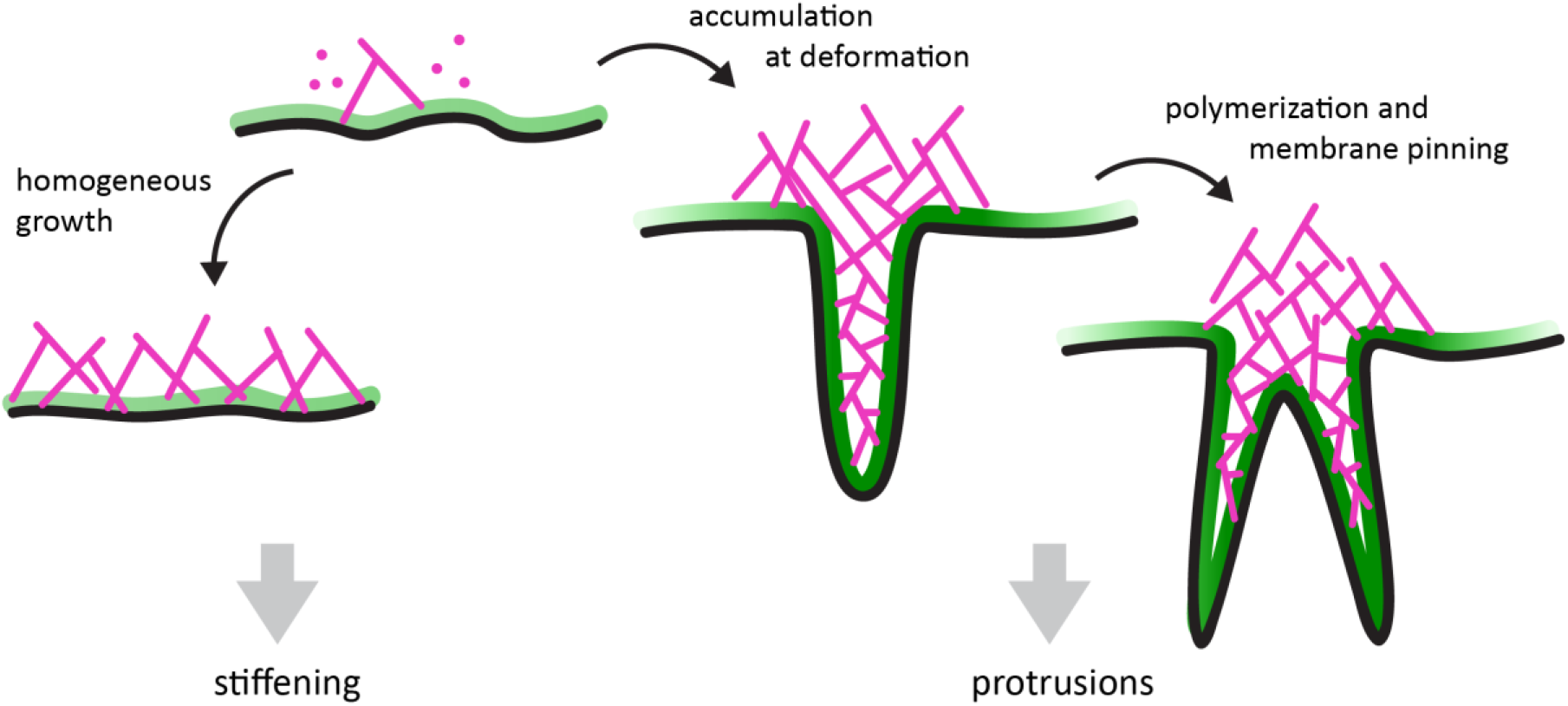
Proposed mechanism of membrane shaping by branched actin cortices. Branched actin networks (magenta) grow on the GUV membrane (black) mediated by VCA (green). Where a sufficient number of growing actin filaments, and thus polymerization forces, accumulate concurrently with a thermally induced membrane deformation, finger-like protrusions can be initiated (middle). The resulting accumulation of yet more actin, and also VCA that mediates actin-membrane-binding, can lead to the formation of more protrusions from the same place, producing a bouquet (right). Otherwise, the membrane is homogeneously coated with a branched actin cortex that mechanically stabilizes the vesicle surface (left).

Arp2/3 and SPIN90-nucleated actin filaments (59), which interpenetrate with branched networks in the cortex of animal cells (2, 3). Finally it will be interesting to include cell-substrate-interactions, which are known to be intimately involved in protrusion formation and maintenance in cells (61).

## Materials and Methods

### Materials

All lipids were purchased from Avanti Polar Lipids and stored in chloroform under argon at -20° C. n-decane (99% pure) was purchased from Arcos Organics. All other chemicals were purchased from Sigma-Aldrich. Chloroform, mineral oil and silicone oil were stored in a glove box with an ambient humidity below 1%.

Human Arp2/3 isoforms ArpC1B/C5L and ArpC1A/C5 were prepared by cloning the appropriate subunits in baculoviruses, and expressing and purifying the protein complexes in Sf21 insect cells. Full details are given in the extended methods in the Supporting Information. Lyophilized rabbit skeletal muscle actin was purchased from Hypermol EK and stored as a 23.8 µM solution in G-buffer at -80°C (see ref. (25) for details). Monomeric (G)-actin was fluorescently labeled with AlexaFluor 488 carboxylic acid succimidyl ester (Invitrogen) (62). The 10xHis-tagged VCA-domain of murine N-WASP (amino acids 400-501) was purified from E. coli BL21 (DE3) cells (44) and fluorescently labeled with AlexaFluor C5 maleimide (Molecular Probes) following the supplier’s protocol. The plasmid was a kind gift from Kristina Ganzinger (AMOLF). Lyophilized Arp2/3 protein complex from porcine brain was purchased from Hypermol EK and stored as a 2.23 µM solution in Arp2/3 storage buffer (20 mM 3-(N-orpholino)propanesulfonic acid (MOPS) pH 7.0, 100 mM KCl, 2 mM MgCl_2_, 5 mM ethylene glycol-bis(β-aminoethyl ether)-tetraacetic acid (EGTA), 1 mM ethylenediaminetetraacetic acid (EDTA), 0.5 mM DL-dithiothreitol (DTT), 0.2 mM adenosine 5’-triphosphate (MgATP), 5% (v/v) glycerol) at -80°C. Thawed aliquots were kept on ice and used within 2 days.

### GUV preparation

GUVs were prepared by eDICE as described in (25), encapsulating 8 μM actin of which 10 % was fluorescently labeled, 50 nM Arp2/3 (human ArpC1B/C5L unless otherwise specified), and varying concentrations of VCA (6.5 μM unless otherwise specified). The inner buffer contained 20 mM Tris(hydroxy-methyl)aminomethane hydrochloride (Tris-HCl) pH 7.4, 50 mM KCl, 2 mM MgCl_2_, 1 mM DTT and 0.5 mM MgATP, and was supplemented with 6.5 % (v/v) Optiprep (Cat. # D1556-250ML) to promote GUV formation, and 0.5 μM proto-catechuate-3,4-dioxygenase (PCD) and 10 mM proto-catechuic acid (PCA) to minimize photobleaching (63). The GUVs were formed in an outer aqueous solution of 190 mM glucose and next diluted into a final buffer containing 170 mM glucose and 10 mM Tris-HCl pH 7.4. To deflate the GUVs, we imposed a slight mismatch between the osmolarity of the inner buffer (168 mOsm/kg) versus outer buffer (182 mOsm/kg), as measured by a freezing point osmometer (Osmomat 010, Gonotec). Lipids were first mixed in chloroform at a molar ratio of 94.985 % 1,2-dioleoyl-sn-glycero-3-phosphocholine (DOPC) : 0.1 % 1,2-distearoyl-sn-glycero-3-phosphoethanolamine-N-[methoxy (polyethylene glycol)-2000] (DOPE-PEG2000) : 5 % 1,2-dioleoyl-sn-glycero-3-[(N-(5-amino-1-carboxypentyl) iminodiacetic acid)succinyl] (DGS-NTA(Ni)): 0.005 % 1,2-dioleoyl-sn-glycero-3-phosphoethanolamine-N-(Cyanine 5) (DOPE-Cy5) and dried in glass vials. In a glovebox, the lipids were dissolved in 50 μL of anhydrous chloroform, diluted with 415 μL n-decane, and then dispersed in a mixture of silicone oil and mineral oil at a 5.3:1.2 volumetric ratio by 15 min sonication on ice. Each GUV preparation was based on 6.5 mL of oil phase containing a final lipid concentration of 0.26 mM. GUVs were prepared within 30 min by manually emulsifying 25 μL of inner aqueous solution with 1 mL of oil phase, layering the emulsion atop 5 mL oil phase and 700 μL outer aqueous solution in a custom-built chamber (64), and centrifuging for 3 minutes.

### Confocal imaging

GUVs were imaged on an inverted Olympus IX81 spinning disk microscope using 491 and 640 nm CW lasers and a 100x oil immersion objective, or on a Leica Stellaris 8 point scanning microscope using a white light laser and a 63x glycerol immersion objective. See Extended Methods for details.

### FRAP experiments

FRAP experiments were performed on a Leica Stellaris 8 confocal microscope (details in the Extended Methods and Table S3 in the Supplementary Information). A small rectangular region (typically 3 × 6.4 μm) centered on the cortex in an equatorial slice of a GUV was photobleached with an intense laser. The fluorescence recovery was extracted by semi-automated image analysis with Fiji (65) and analyzed with a custom written python script as explained in Fig. S3.

### Photoablation

GUV cortices were ablated in their entirety or at the base of membrane spikes by illumination with a 1.6 W UV laser diode (405 nm) on the Leica Stellaris 8 confocal microscope. Before and after ablation, a z-stack was recorded to capture the GUV shapes. In case of whole-cortex ablation, rounding of the GUV surface and the return of thermal membrane undulations was already visible after 3 ablation rounds. In case of local ablation of spikes, a rectangular region of a size matching the spike’s dimensions (typically ∼ 3 × 1.5 μm) was illuminated with the laser diode at full power for several consecutive rounds of ablation (see Table S3). We checked by visual inspection of the actin and membrane signals that no membrane damage occurred.

### Tracking membrane displacements

Time-lapse videos of the equatorial slices of two GUVs (50 s at 1 fps) were visualized in a color-coded projection using the Fiji plugin Temporal Color Code (66). The average membrane displacements were measured by analyzing the same videos with a custom-written python script kindly provided by Lennard van Buren. The images were first smoothed with a 3 px 2D Gaussian filter. Radial line intensities were extracted in 1° angular intervals and smoothed by a 3 px 1D Gaussian filter. The membrane position was set as the location of maximum intensity. Displacements of each angular membrane segment between consecutive frames were measured, and the root mean square displacement X_rms_ of the ensemble of membrane segments was computed.

### Analysis of GUV sizes and protrusions

Sizes of GUVs and of membrane protrusions were manually analyzed in Fiji (65). GUV sizes were measured by fitting the GUV with the largest circle around the body (excluding membrane protrusions). Protrusion widths were determined from line profiles drawn across the protrusion base as the distance between the membrane intensity peaks on either side. If the protrusion base was so thin that no separate peaks could be resolved, we set the width to zero and classified the protrusion as a tube, otherwise we classified it as a spike. Protrusion lengths were measured from the base to the outermost visible tip. The number of protrusions per bouquet was counted manually, and the widest angle between any two protrusions in the same bouquet was measured using the Fiji ‘Angle’ tool. Actin intensity profiles were extracted by manually tracing the GUV membrane with an 8 px-wide segmented line ROI. To achieve unbiased classification of protrusion prevalence, we first inspected only the membrane channel in a z-stack of each GUV, and noted whether any membrane structures (tubes, spikes, bouquets, inward-pointing fuzz) were visible. Afterwards, we inspected the actin channel for any visible actin enrichment at any of the membrane structures. Quantitative comparisons for Fig. 4 were done on samples produced on the same day, to exclude effects from day-to-day variations in encapsulation efficiency. We confirmed separately that protrusions were more prevalent in GUVs containing 2.6 uM VCA than either 0.65 or 6.5 uM VCA (N=2 days for each comparison).

### Actin polymerization assay

Actin polymerization speeds in bulk solution were quantified spectrophotometrically by pyrene assays (67, 68). We measured actin polymerization at 4 μM actin, 650 nM VCA and 50 nM Arp2/3, in the same buffer as the inner aqueous solution of the GUVs. Details are given in the extended methods in the Supporting Information.

## Code availability

Python-based data analysis scripts used in this work are available on GitHub: github.com/BioSoftMatterGroup/GUV-deformations

## Acknowledgments

The GoldenBac vectors and guidance were kindly provided by Thomas Puehringer in the Alessandro Costa lab at the Francis Crick Institute. We thank Kristina Ganzinger (AMOLF) for the kind gift of the VCA plasmid and for assistance with pyrene assays, Lennard van Buren for the membrane tracking software, Jeffrey den Haan and Erik van Lagen for protein purification, James Conboy for assistance with the cytochalasin D experiments, and Jérémie Capoulade and Heike Glauner (Leica) for help with FCS measurements. We thank Dyche Mullins (UCSF), Stephan Grill (MPI-CBG Dresden), Dan Fletcher (UC Berkeley) and Guillaume Charras (UCL) for helpful discussions about this work. We acknowledge financial support from The Netherlands Organization of Scientific Research (NWO/OCW) Gravitation program Building A Synthetic Cell (BaSyC) (024.003.019) and the Kavli Synergy program of the Kavli Institute of Nanoscience Delft (to FF). MM and MW were supported by the Francis Crick Institute, which receives its core funding from Cancer Research UK (CC2096), the UK Medical Research Council (CC2096), and the Wellcome Trust (CC2096) as well as by the European Research Council (ERC) under the European Union’s Horizon 2020 research and innovation programme (grant agreement No 810207 to MW).

## Notes

https://github.com/BioSoftMatterGroup/GUV-deformations

### Competing Interest Statement

The authors have declared no competing interest.

## References

1. P. Chugh, E. K. Paluch, The actin cortex at a glance. J. Cell Sci. 131, jcs186254 (2018).

2. M. Bovellan, et al., Cellular control of cortical actin nucleation. Curr. Biol. 24, 1628–1635 (2014).

3. M. Fritzsche, C. Erlenkämper, E. Moeendarbary, G. Charras, K. Kruse, Actin kinetics shapes cortical network structure and mechanics. Sci. Adv. 2, e1501337 (2016).

4. Rohatgi R., et al., The interaction between N-WASP and the Arp2/3 complex links Cdc42-dependent signals to actin assembly. Mol. Biol. Cell 97, 221–231 (1999).

5. A. M. Gautreau, F. E. Fregoso, G. Simanov, R. Dominguez, Nucleation, stabilization, and disassembly of branched actin networks. Trends Cell Biol. 32, 421–432 (2022).

6. M. Kelkar, P. Bohec, G. Charras, Mechanics of the cellular actin cortex: From signalling to shape change. Curr. Opin. Cell Biol. 66, 69–78 (2020).

7. M. Fritzsche, A. Lewalle, T. Duke, K. Kruse, G. Charras, Analysis of turnover dynamics of the submembranous actin cortex. Mol. Biol. Cell 24, 757–767 (2013).

8. N. Vadnjal, et al., Proteomic analysis of the actin cortex in interphase and mitosis. J. Cell Sci. 135, jcs259993 (2022).

9. M. Biro, et al., Cell cortex composition and homeostasis resolved by integrating proteomics and quantitative imaging. Cytoskeleton 70, 741–754 (2013).

10. A. Mogilner, B. Rubinstein, The physics of filopodial protrusion. Biophys. J. 89, 782–795 (2005).

11. D. Vignjevic, et al., Role of fascin in filopodial protrusion. J. Cell Biol. 174, 863–875 (2006).

12. G. Sekerková, L. Zheng, P. A. Loomis, E. Mugnaini, J. R. Bartles, Espins and the actin cytoskeleton of hair cell stereocilia and sensory cell microvilli. Cell. Mol. Life Sci. 63, 2329–2341 (2006).

13. F.-C. Tsai, G. H. Koenderink, Shape control of lipid bilayer membranes by confined actin bundles. Soft Matter 11, 8834–8847 (2015).

14. Y. Bashirzadeh, H. Moghimianavval, A. P. Liu, Encapsulated actomyosin patterns drive cell-like membrane shape changes. iScience 25, 104236 (2022).

15. T. M. Svitkina, et al., Mechanism of filopodia initiation by reorganization of a dendritic network. J. Cell Biol. 160, 409–421 (2003).

16. C. Yang, et al., Novel roles of formin mDia2 in lamellipodia and filopodia formation in motile cells. PLoS Biol. 5, 2624–2645 (2007).

17. A. B. Bohil, B. W. Robertson, R. E. Cheney, Myosin-X is a molecular motor that functions in filopodia formation. PNAS 103, 12411–12416 (2006).

18. C. Simon, et al., Actin dynamics drive cell-like membrane deformation. Nat. Phys. 15, 602–609 (2019).

19. G. Salbreux, G. T. Charras, E. K. Paluch, Actin cortex mechanics and cellular morphogenesis. Trends Cell Biol. 22, 536–545 (2012).

20. K. Carvalho, et al., Cell-sized liposomes reveal how actomyosin cortical tension drives shape change. PNAS 110, 16456–16461 (2013).

21. E. Loiseau, et al., Shape remodeling and blebbing of active cytoskeletal vesicles. Sci. Adv. 2, e1500465 (2016).

22. T. Litschel, et al., Reconstitution of contractile actomyosin rings in vesicles. Nat. Commun. 12, 2254 (2021).

23. L. L. Pontani, et al., Reconstitution of an actin cortex inside a liposome. Biophys. J. 96, 192–198 (2009).

24. K. Dürre, et al., Capping protein-controlled actin polymerization shapes lipid membranes. Nat. Commun. 9, 1630 (2018).

25. L. Baldauf, F. Frey, M. A. Perez, T. Idema, G. H. Koenderink, Reconstituted branched actin networks sense and generate micron-scale membrane curvature. biorxiv Prepr., 2022.08.31.505969 (2022).

26. S. Suetsugu, Activation of nucleation promoting factors for directional actin filament elongation: Allosteric regulation and multimerization on the membrane. Semin. Cell Dev. Biol. 24, 267–271 (2013).

27. P. Echave, I. J. Conlon, A. C. Lloyd, Cell size regulation in mammalian cells. Cell Cycle 6, 218–224 (2007).

28. P. Chugh, et al., Actin cortex architecture regulates cell surface tension. Nat. Cell Biol. 19, 689–697 (2017).

29. M. P. Clausen, H. Colin-York, F. Schneider, C. Eggeling, M. Fritzsche, Dissecting the actin cortex density and membrane-cortex distance in living cells by super-resolution microscopy. J. Phys. D. Appl. Phys. 50, 064002 (2017).

30. V. Laplaud, et al., Pinching the cortex of live cells reveals thickness instabilities caused by myosin II motors. Sci. Adv. 7, eabe3640 (2021).

31. M. P. Murrell, et al., Spreading dynamics of biomimetic actin cortices. Biophys. J. 100, 1400–1409 (2011).

32. D. Raz-Ben Aroush, et al., Actin turnover in lamellipodial fragments. Curr. Biol. 27, 2963– 2973 (2017).

33. D. J. Barry, C. H. Durkin, J. V. Abella, M. Way, Open source software for quantification of cell migration, protrusions, and fluorescence intensities. J. Cell Biol. 209, 163–180 (2015).

34. G. Romet-Lemonne, A. Jégou, The dynamic instability of actin filament barbed ends. J. Cell Biol. 220, e202102020 (2021).

35. M. M. A. E. Claessens, F. A. M. Leermakers, F. A. Hoekstra, M. A. C. Stuart, Osmotic shrinkage and reswelling of giant vesicles composed of dioleoylphosphatidylglycerol and cholesterol. Biochim. Biophys. Acta 1778, 890–895 (2008).

36. U. Seifert, K. Berndl, R. Lipowsky, Shape transformation of vesicles: Phase diagram for spontaneous-curvature and bilayer-coupling models. Phys. Rev. A 44, 1182–1202 (1991).

37. M. Mayer, M. Depken, J. S. Bois, F. Jülicher, S. W. Grill, Anisotropies in cortical tension reveal the physical basis of polarizing cortical flows. Nature 467, 617–621 (2010).

38. C. Guillot, T. Lecuit, Adhesion disengagement uncouples intrinsic and extrinsic forces to drive cytokinesis in epithelial tissues. Dev. Cell 24, 227–241 (2013).

39. N. S. Gov, Dynamics and morphology of microvilli driven by actin polymerization. Phys. Rev. Lett. 97, 018101 (2006).

40. N. S. Gov, A. Gopinathan, Dynamics of membranes driven by actin polymerization. Biophys. J. 90, 454–469 (2006).

41. R. D. Mullins, P. Bieling, D. A. Fletcher, From solution to surface to filament: actin flux into branched networks. Biophys. Rev. 10, 1537–1551 (2018).

42. S. Ghosh, S. Gutti, D. Chaudhuri, Pattern formation, pulsation and traveling wave on active spherical membranes. Soft Matter 47, 10583–10778 (2021).

43. J. V. G. Abella, et al., Isoform diversity in the Arp2/3 complex determines actin filament dynamics. Nat. Cell Biol. 18, 76–86 (2016).

44. Sonal, et al., Myosin-II activity generates a dynamic steady state with continuous actin turnover in a minimal actin cortex. J. Cell Sci. 132, jcs299899 (2019).

45. C. Co, D. T. Wong, S. Gierke, V. Chang, J. Taunton, Mechanism of actin network attachment to moving membranes: Barbed end capture by N-WASP WH2 domains. Cell 128, 901–913 (2007).

46. I. Weisswange, T. P. Newsome, S. Schleich, M. Way, The rate of N-WASP exchange limits the extent of ARP2/3-complex-dependent actin-based motility. Nature 458, 87–91 (2009).

47. N. Selve, A. Wegner, Rate of treadmilling of actin filaments in vitro. J. Mol. Biol. 187, 627– 631 (1986).

48. A. E. Carlsson, The effect of branching on the critical concentration and average filament length of actin. Biophys. J. 89, 130–140 (2005).

49. J. Rosenblatt, B. J. Agnew, H. Abe, J. R. Bamburg, T. J. Mitchison, Xenopus actin depolymerizing factor/cofilin (XAC) is responsible for the turnover of actin filaments in Listeria monocytogenes tails. J. Cell Biol. 136, 1323–1332 (1997).

50. N. G. Pandit, et al., Force and phosphate release from Arp2/3 complex promote dissociation of actin filament branches. PNAS 117, 13519–13528 (2020).

51. A. M. Weaver, et al., Cortactin promotes and stabilizes Arp2/3-induced actin filament network formation. Curr. Biol. 11, 370–374 (2001).

52. C. Galloni, et al., MICAL2 enhances branched actin network disassembly by oxidizing arp3b-containing arp2/3 complexes. J. Cell Biol. 220, e202102043 (2021).

53. L. Song, et al., Structure of staphylococcal a-hemolysin, a heptameric transmembrane pore. Science (80-.). 274, 1859–1866 (1996).

54. A. Rørvig-Lund, A. Bahadori, S. Semsey, P. M. Bendix, L. B. Oddershede, Vesicle fusion triggered by optically heated gold nanoparticles. Nano Lett. 15, 4183–4188 (2015).

55. N. Leijnse, et al., Filopodia rotate and coil by actively generating twist in their actin shaft. Nat. Commun. 13, 1636 (2022).

56. A. P. Liu, et al., Membrane-induced bundling of actinfilaments. Nat. Phys. 4, 789–793 (2008).

57. J. Weichsel, P. L. Geissler, The more the tubular: Dynamic bundling of actin filaments for membrane tube formation. PLoS Comput. Biol. 12, e1004982 (2016).

58. P. Bieling, et al., WH2 and proline-rich domains of WASP-family proteins collaborate to accelerate actin filament elongation. EMBO J., e201797039 (2017).

59. L. Cao, F. Ghasemi, M. Way, A. Jégou, G. Romet-Lemonne, Nucleation and stability of branched versus linear Arp2/3-generated actin filaments. biorxiv Prepr. (2022) https://doi.org/10.1101/2022.05.06.490861.

60. A. Y. Pollitt, R. H. Insall, WASP and SCAR/WAVE proteins: The drivers of actin assembly. J. Cell Sci. 122, 2575–2578 (2009).

61. P. Sens, Stick-slip model for actin-driven cell protrusions, cell polarization, and crawling. PNAS 117, 24670–24678 (2020).

62. J. Alvarado, G. H. Koenderink, “Reconstituting cytoskeletal contraction events with biomimetic actin-myosin active gels” in Methods in Cell Biology, (Elsevier Inc., 2015), pp. 83–103.

63. C. E. Aitken, R. A. Marshall, J. D. Puglisi, An oxygen scavenging system for improvement of dye stability in single-molecule fluorescence experiments. Biophys. J. 94, 1826–1835 (2008).

64. L. Van De Cauter, et al., Optimized cDICE for efficient reconstitution of biological systems in giant unilamellar vesicles. ACS Synth. Biol. 10, 1690–1702 (2021).

65. J. Schindelin, et al., Fiji: An open-source platform for biological-image analysis. Nat. Methods 9, 676–682 (2012).

66. K. Miura, J. Schindelin, Temporal Colorcode (2018).

67. J. A. Cooper, S. B. Walker, T. D. Pollard, Pyrene actin: documentation of the validity of a sensitive assay for actin polymerization. J. Muscle Res. Cell Motil. 4, 253–262 (1983).

68. L. K. Doolittle, M. K. Rosen, S. B. Padrick, “Measurement and analysis of in vitro actin polymerization” in Methods in Molecular Biology, A. S. Coutts, Ed. (Springer Science and Business Media LLC, 2013), pp. 273–293.

