## Supporting Information for "Biomimetic actin cortices shape cell-sized lipid vesicles"

[b] Present address: Institute of Science and Technology Austria, 3400 Klosterneuburg, Austria

**This PDF file includes:**

Supporting text:
  Modeling the formation of membrane protrusions
  Extended methods
Figures S1 to S15
Tables S1 to S3
Legends for Movies S1 to S3
SI References

**Other supporting materials for this manuscript include the following:**

Movies S1 to S3
Software:
https://github.com/BioSoftMatterGroup/GUV-deformations
https://gitlab.tudelft.nl/biosoft/personal/vesicle-fluctuation-analysis

**Supporting Information Text**

**Modelling the formation of membrane protrusions**

To model membrane deformation as a function of the density of the Arp2/3 activator VCA on the membrane, we assumed a VCA response function *f(n)* that describes how the VCA density *n* affects actin polymerization. Multiplying *f(n)* by the (maximum) actin polymerization velocity *v* gives the effective actin polymerization velocity, which determines how fast the membrane is pushed outward. We chose a sigmoidal shape for *f(n)* to reflect the reasonable assumptions that we need a minimal concentration of VCA for actin to start polymerizing, that actin polymerization saturates at high VCA density and that in between we expect actin polymerization to increase linearly with VCA density

$$f(n) = \frac{1}{2}\left[\tanh\left(\mathrm{a}\,(\mathrm{n}-\mathrm{b})\right)+1\right]. \tag{S1}$$

We set $b$ = 0.5, i.e., the inflection point of the response function is at half the membrane coverage of VCA. We set $a = 10$ to interpolate smoothly between the regime of low and high VCA density. The resulting function is plotted in Fig. S8. We note that our qualitative results did not depend on the exact choice of *a* and *b* (Fig. S9).

We consider the VCA density *n* on a 2D membrane. We use the Monge gauge to describe the membrane shape, i.e., we use the membrane height field *h(x,y),* the distance between the membrane position and some flat reference plane. Because we limited our theory to the initiation of membrane protrusions, we can work in the limit of small deformations, and rewrote the total membrane curvature in Eq. (1) of the main text as $K^2 \approx (\nabla^2 h)^2$. In addition, we assumed rotational symmetry, i.e., $n = n(r, t)$ and $h = h(r, t)$, and rewrote Eqs. (1) and (2) of the main text in polar coordinates as a function of the distance *r* to the z-axis,

$$\begin{aligned}\frac{\partial n(r,t)}{\partial t} &= D\left(\frac{1}{r}\frac{\partial}{\partial r}+\frac{\partial^2}{\partial r^2}\right)n(r,t)\\ &\quad + \alpha\, n(r,t)\big(1-n(r,t)\big)\left[\left(\frac{1}{r}\frac{\partial}{\partial r}+\frac{\partial^2}{\partial r^2}\right)h(r,t)\right]^2\end{aligned} \tag{S2}$$

$$\partial_t h(r,t) = v\, f\big(n(r,t)\big) - w\, h(r,t) \tag{S3}$$

with the Laplacian in polar coordinates, $\nabla^2 = \left(\frac{1}{\mathrm{r}}\frac{\partial}{\partial \mathrm{r}}+\frac{\partial^2}{\partial \mathrm{r}^2}\right)$, the diffusion constant $D$ = 1 $\mu m^2/s$ (1), the polymerization velocity of actin $v$ = 1 $\mu m s^{-1}$ (12), the mechanical membrane relaxation rate $w$ = 0.013$s^{-1}$ (11) and the enrichment constant $\alpha$ = 0.1 $\mu m^2/s$. We note that our results remained qualitatively similar even when we decreased $\alpha$ by an order of magnitude (Fig. S10).

To solve Eqs. (S2) and (S3), we had to set up suitable initial conditions. To represent the cases of a low, intermediate and high homogeneous initial density of VCA, we assumed that the membrane coverage was sparse, i.e., $n_l(r, 0)=0.1$, intermediate, i.e., $n_i(r, 0)=0.5$ or dense, i.e., $n_h(r, 0)=0.9$. To represent a localized membrane deformation, we chose the membrane height function to initially have a Gaussian shape, $h(r,0) = A\exp(-\frac{r^2}{\sigma^2})$ with amplitude $A$ and width $\sigma$. As we wanted to investigate the effect of a small initial membrane perturbation, we chose $A = 0.1$ µm and $\sigma = 0.2$ µm, similar to (2).

To complete the model, we also had to specify the boundary conditions. To shorten our notation, in the following we use primes to denote radial derivatives. We considered a membrane within a spherical domain. To avoid numerical instability, we chose a cutoff of $r_{min} = 10^{-6}$ µm at the inner domain boundary. We set the outer domain boundary to $r_{\max} =$ 10 µm. We assumed a no-flux boundary condition of VCA at both boundaries, i.e. $n'(r_{min}, t) = 0$ and $n'(r_{max}, t) = 0$, respectively. Moreover, we assumed that membrane protrusions are pushed outward by more than one actin filament sharing the membrane load, as shown in simulations (3). Therefore, we assumed that the membrane remains flat at the protrusion center, $h'(r_{min}, t) = 0$. Finally, we assumed that the membrane also remains flat far from the initial deformation, $h'(r_{max}, t) = 0$. We solved Eqs. (S2) and (S3) in *Mathematica* with the numerical method “Method-Of-Lines”. For the spatial discretization we used the ‘TensorProductGrid’ method. In order to provide sufficient numerical accuracy, we manually set the minimum number of points for each dimension of the grid to 1000, by using the ‘MinPoints’ method. To represent the initial time domain of membrane protrusion initiation, we chose the time range between 0 and 20 s. The parameter values that were used for the calculations are summarized in Table S1.

### Extended methods

**Expression and purification of human Arp2/3 isoforms: Cloning and construction of baculoviruses.** The open reading frames (ORFs) for all Arp2/3 complex subunits were synthesized with codon-optimization for expression in insect cells by Twist Bioscience. The ArpC3 ORF was synthesized with an additional in frame TEV cleavage site followed by a Twin-Strep-tag at its C-terminus. The seven Arp2/3 complex subunit ORFs were amplified from the Twist Bioscience templates with 15 bp overhangs compatible with the appropriate termini of the linearized pGB vectors (4), which were also amplified at the same time using compatible primers with Q5 polymerase (New England Biolabs). The PCR products and linearized, amplified pGB vectors (ThermoFisher Scientific) were treated with Dpn I restriction enzyme (ThermoFisher Scientific) for 1 hour at 37° C prior to gel purification (QIAquick Gel Extraction Kit, Qiagen). Individual purified Arp2/3 subunit ORFs were mixed with the corresponding pGB vector (2:1 insert:vector) in a 10 µL In-Fusion reaction (Takara Biosciences) and incubated for 1 h at 50° C. Chemically competent NEB 5-alpha Competent cells (New England Biolabs) were transformed with 5 µL of the In-fusion reaction and plated on gentamicin-containing LB agar plates. The DNA sequence of the inserted genes and flanking BsaI sites from single clones were verified by Sanger sequencing. The Arp2/3 complex subunit ORFs were then released from the pGB vectors by cleavage with the BsaI-HF v2 restriction enzyme (New England Biolabs) and simultaneously ligated into the final destination vector pGBDest, based on the inter-compatible overhangs flanking the ORFs in a Golden Gate reaction following the method of (4). In addition, the final position in the co-expression construct was closed off by a “dummy” pGB-dummy-08-16 (4) (BCCM/GeneCorner) that was inserted between the end of the 7th ORF and the pGBDest vector. The ligation mixture was transformed into chemically competent NEB 5-alpha Competent cells (New England Biolabs) and clones were selected on kanamycin containing LB agar plates. The final expression vector containing the seven Arp2/3 subunit ORFs was verified by Sanger sequencing before being transformed into MAX Efficiency DH10Bac Competent E. coli (Invitrogen) on to antibiotic plates with blue/white X-Gal/IPTG screening. White colonies were used to inoculate 3 ml cultures for overnight growth, followed by purification of bacmid DNA using QIAprep Spin Miniprep kit (Qiagen).

**Expression and purification of human Arp2/3 isoforms: Baculovirus and protein production.** To generate baculoviruses, bacmid DNA (2 µg) was mixed with 100 µL SF900-III medium (Life Technologies) and 3 µL FuGene HD transfection reagent, and incubated for 15 min before being added dropwise to 1 x $10^6$ adherent Sf21 insect cells in a 6-well plate with 2 mL of SF900-III medium at 27°C. After 3 days, the supernatant (P1 virus) was added to a 50 mL culture of Sf21 cells (1–2x$10^6$ cells/mL) in SF900-III medium with constant shaking at 110 rpm at 27°C. After 3 days, 50 µL of supernatant (P2 virus) was used to infect a second 50 mL Sf21 culture and incubated for 3 days. The resulting supernatant (P3 virus) was stored at 4°C and 500 µL were used to infect 0.5 L of Sf21 insect cells at 1–2x$10^6$ cells/mL for protein production. Three days after infection, cell pellets were harvested by centrifugation, washed with phosphate buffer saline (PBS), flash frozen in liquid nitrogen, and stored at -80°C. Frozen cell pellets were resuspended to a final volume of 50 mL in purification buffer (50 mM Tris pH 8, 150 mM NaCl, 2 mM MgCl2, 2.5 % v/v glycerol, 1 mM DTT, 0.2 mM Mg-ATP) supplemented with cOmplete EDTA-free Protease Inhibitor Cocktail (Roche) and BaseMuncher Endonuclease (5 µL per 50 mL) (Abcam, Cat. # ab270049). Resuspended cells were lysed by sonication for 120 seconds (5 s on followed by 10 s off) at 4° C using a Branson Digital Sonifier at 40 % power. After 1 hour incubation, EDTA and EGTA were added to a final concentration of 1 mM and 5 mM respectively. The lysate was then clarified by ultracentrifugation in a Type 45 Ti rotor (Beckman) at 41,000 rpm for 45 min, followed by passage through a 0.45 µm syringe filter. The resulting filtrate was loaded onto a 1 mL StrepTrap XT column (Cytiva) attached to an ÄKTApure system (GE Healthcare) at a rate of 1 mL/min. After washing with 30 column volumes of purification buffer, the bound Arp2/3 complex was eluted with 15 column volumes of Buffer BXT (IBA) supplemented with 2 mM ATP, 1 mM DTT, 5 mM EGTA, 2 mM $MgCl_2$ and 2.5 % v/v glycerol. Fractions containing Arp2/3 were pooled and loaded on a Superdex 200 Increase 10/300 GL column that was pre-equilibrated with purification buffer containing 5 % v/v glycerol. Column fractions (0.4 mL) were analyzed by SDS-PAGE (4 – 12% Bolt Bis-Tris Plus gels (Invitrogen) stained with Quick Coomassie Stain (Neo Biotech). Fractions with Arp2/3 complexes were analyzed by mass photometry (Refeyn) to confirm that all seven subunits were stoichiometric prior to being pooled and concentrated to ~4-5 mg/mL. Small aliquots were flash-frozen in liquid $N_2$ and stored at -80° C. An SDS PAGE gel of Arp2/3C1A/C5 and Arp2/3C1B/C5L is shown in Fig. S15.

**Microscopy.** GUVs were observed in ibidi 18-well µ-slides or in chambers made of # 1.5 coverslips (Superior Marienfeld) separated by silicone spacers (Sigma Aldrich) previously passivated by incubating for 15 min with a 0.1 mg/mL β-casein solution in 10 mM Tris-HCl pH 7.4, followed by rinsing with MilliQ water and drying under $N_2$ flow. Upon GUV addition, the chambers were closed to prevent solvent evaporation and maintain constant osmotic conditions. Images were acquired on an inverted Olympus IX81 confocal spinning disk microscope using 491 and 640 nm CW lasers, a 100x oil immersion objective (UPlanSApo, WD 0.13 mm, NA 1.4) and an EM-CCD Andor iXon X3 DU897 camera, or on an inverted Leica Stellaris 8 point scanning confocal microscope using a white light laser, a 63x glycerol immersion objective (HC PL APO, WD 0.3 mm, NA 1.2) and HyD detectors operated in photon counting mode (see imaging settings see Table S2).

**Fluorescence Correlation Spectroscopy.** We used fluorescence correlation spectroscopy (FCS) to determine the diffusion coefficient of fluorescently labeled actin monomers in our GUVs, and assess whether there was any free dye present in the

samples. All measurements were performed on a Leica Stellaris DMI8 microscope equipped with a white light laser and a 63x water immersion objective (HC Plan APO 63x/1.20 W Corr CS2), at a sample temperature of 25° C (controlled by an Okolab environmental control box). We measured on vesicles in # 1.5 ibidi 18-well μ-slides, 5.0 μm above the coverslip surface. Confocal volume and structure parameter of the laser beam were calibrated each day, fitting decorrelation curves extracted from 60 s long fluorescence traces using the known diffusion constant of 10 nM free AlexaFluor488 dye in MilliQ water (414 $\mu m^2/s$ (5)). All other measurements were performed in G-buffer (5 mM Tris-HCl pH 7.8, 0.2 mM $CaCl_2$, 1 mM DTT, 0.2 mM NaATP) supplemented with 6.5 % Optiprep, to reflect the buffer conditions inside the GUVs while keeping actin monomeric. Photon yields were optimized separately each day on reference samples of 10 nM free dye in the final buffer. Optimal imaging conditions typically required a laser intensity of 1.2 % at 491 nm on the white light laser.
We first measured the diffusion time of 10 nM free dye in the final buffer, which we extracted from 60 s fluorescence traces. The autocorrelation curves were fitted assuming a single fluorescent species undergoing normal diffusion with a constant diffusion coefficient, and including a triplet contribution (6):

$$G(\tau) = \left[1 - \tau + e^{-\frac{t}{\tau_T}}\right] \cdot \frac{1}{N(1-\tau)\left(1+\frac{\tau}{\tau_D}\right)} \cdot \left[1 + \left(\frac{\tau}{\tau_D}\right)\left(\frac{\omega}{\omega_2}\right)^2\right]^{-1/2}$$

Here, $\tau$ is the lag time, $\tau_T$ and $\tau_D$ are the characteristic times of the fluorophore's triplet state and the molecular diffusion respectively, $\frac{\omega}{\omega_2}$ is the structure parameter, and $N=cN_AV_0$ is the typical number of fluorescent particles in the confocal volume $V_0$. All fitting was performed using the commercial Leica LAS X FCS software. To measure G-actin diffusion coefficients and the relative contribution of G-actin and free dye, we performed the same measurements on solutions of monomeric (G-)actin with a nominal concentration of 10 nM AlexaFluor488-labeled monomers. Here, the autocorrelation curves showed a bimodal decay due to diffusion, where the shorter characteristic timescale was associated with the diffusion of the free dye molecules, and the longer timescale was associated with diffusion of labeled actin monomers. We fitted the curves using the same model as in case of the free dye, but with two diffusing species. One characteristic diffusion time was fixed at the value measured for the free dye alone, and the other was left as a free parameter. To ensure stable fitting, we constrained triplet times to less than 5 μs and triplet fractions to below 25 %. This analysis revealed that free dye contributed significantly to overall `actin' fluorescence (Fig. S1 A) and yielded an actin monomer diffusion coefficient of 96.7 ± 8.4 $\mu m^2/s$ (Fig. S1 B). Note that G-actin diffusion was slowed by over 50 % in the buffer with Optiprep compared to F-buffer, even though the buffer viscosity increased by only 2 % upon adding 6.5 % Optiprep (Fig. S1 C, D). The diffusion coefficient of free AlexaFluor488 dye molecules was much less affected by the presence of Optiprep, suggesting that Optiprep associates with G-actin, forming a hydration shell that reduces protein mobility and also slows down actin polymerization (7).

**Viscosity measurements.** Buffer viscosities were measured on a Kinexus Malvern Pro rheometer using a stainless steel cone-plate geometry with 40 mm radius and a 1° angle. We performed a viscometry ramp, measuring viscosity as a function of shear rate from 0.5 $s^{-1}$ to 100 $s^{-1}$ in a logarithmic ramp with 10 samples per decade, with a total ramp time of ~2 min. This measurement was repeated at least three times for every buffer composition, and the mean viscosity was calculated from all values measured above 30 $s^{-1}$. Values at lower shear rates were dominated by the rheometer's inertia and thus excluded.

**FRAP.** We analyzed FRAP data by first extracting an intensity profile along the GUV cortex and smoothing it (Fig. S2 A, B) for each frame. To correct for photobleaching during acquisition, we then defined a reference ROI along with the bleached FRAP ROI (Fig. S2 C, D). As a reference region, we chose a section of at least 10 µm long and no closer than 10 µm to the bleached region, to avoid any artefacts from broadening of the bleached region. The reference region was chosen individually for each GUV to ensure that it did not show large intensity fluctuations over time, which might impact the analysis. Such fluctuations sometimes happened when a bright actin spot in the GUV lumen came close to the cortex. We then normalized the entire profile to the reference intensity (Fig. S2 E) and computed an average intensity in the FRAP region for every frame. To avoid artefacts from spatial inhomogeneities in cortical actin fluorescence, we normalized the data again such that the average intensity in the FRAP region before bleaching ($t<0$) was 1. We computed the asymptotic recovered intensity $I_{inf}$ by calculating the average intensity in the last three time points, and fitted the data to a single exponential decay with characteristic time $\tau$, recovering to that asymptotic value according to $I(t) = I_{inf} - A \cdot exp(-\frac{t}{\tau})$). We considered fits valid if the correlation coefficient was better than $R^2$=0.8.
To assess fluorescence recovery in the membrane, we performed the same procedure on vesicles in which we increased the membrane dye concentration 100-fold from 0.005 to 0.5 % compared to our other experiments, where we kept membrane labeling as low as possible to avoid any artefacts from fluorescence crosstalk into the actin channel. The acquisition and analysis were identical to the actin cortex experiments, but we acquired images with shorter pixel dwell times, to be able to capture the rapid fluorescence recovery in the membrane. Measuring VCA mobility on the membrane again followed the same procedure.

**Pyrene actin polymerization assays.** Pyrene assays to measure the kinetics of actin polymerization were performed and analyzed following the protocol in (8). Actin, VCA and Arp2/3 were used at 4 µM, 0.65 µM and 50 nM, respectively, with 5 % of actin monomers being pyrene-labeled. Before triggering polymerization by mixing actin monomers with the regulatory proteins and salts, we incubated actin in Mg-G-buffer (5 mM Tris-HCl pH 7.4, 0.2 mM $MgCl_2$, 1 mM DTT, 0.2 mM MgATP) for 2 min, to exchange actin-bound calcium ions for magnesium. The measurements were performed in F-buffer (20 mM Tris-HCl pH 7.4, 50 mM KCl, 2 mM $MgCl_2$, 1 mM DTT, 0.5 mM MgATP) supplemented with 6.5 % (v/v) Optiprep, to match the buffer composition inside GUVs. All measurements were performed in 55 µL quartz cuvettes with a 3 mm optical path length (Hellma Analytics) in a Duetta fluorescence and absorbance spectrometer (Horiba Scientific) equipped with an 80 W S/N 1344-DL lamp and TC1 temperature controller (Quantum Northwest) at 25.2 ± 0.2°C. The samples were excited at 365 nm with a 10 nm excitation window, and the emission was recorded at 407 nm with a 5 nm window and a 1 s integration time per time point. Measurements were left running until the fluorescence intensity plateaued. Polymerization speeds were extracted from the slope of the polymerization curve at the point where half of all actin was polymerized (8). At least two independent measurements were performed with each Arp2/3 isoform, and four for spontaneous actin polymerization.

**Actin depolymerization by cytochalasin D.** Exposure of biological membranes to high laser light intensities has been reported to produce membrane shape transformations and can lead to oxidative damage of the sample (9). To independently confirm our conclusion from photoablation assays that removing the actin cortex from GUVs returns their shapes to those predicted for fluid vesicle membranes, we therefore also performed an experiment where we chemically depolymerized actin. To this end, we used the membrane-permeable drug cytochalasin D, which is thought to act specifically on dynamic actin (10, 11). We prepared GUVs with cortices composed of 8 µM actin, 6.5 µMVCA, and 50 nM ArpC1B/C5L, and acquired confocal z-stacks of the sample (Fig. S6 A). As expected, we observed many globally deformed GUVs with bright actin cortices. We then added cytochalasin D to a final concentration of 10 µM based on typical application of the drug in cells (12) and incubated the GUVs with the drug for 90 minutes. Finally, we acquired another set of confocal z-stacks (Fig. S6 B), which revealed that the distinctive actin cortices had disappeared, and most vesicles instead showed a homogeneous, bright cytosolic actin signal. Importantly, vesicles had spherical shapes and we no longer observed any vesicles with stable, large-scale deformations.

**Estimating VCA surface coverage**

To estimate the surface density of VCA in GUVs, we assumed that a typical GUV is spherical and has a radius of $r_{GUV} = 4.5\mu m$ equal to the mean radius (7). Further, we assumed that VCA is a globular protein with a molecular mass of 15 kDa and the average protein density of 1.44 g/cm$^3$ (13), and that the VCA encapsulation efficiency is 100%. The surface coverage $S$ is then

$$S = \frac{A_{VCA}^{total}}{A_{GUV}} \quad \text{(S4)}$$

where $A_{VCA}^{total}$ is the total surface area taken up by VCA and $A_{GUV}$ is the surface area of a GUV. The total area of VCA is the product of the total number of VCA molecules in a typical GUV ($N_{VCA}$) and the cross-sectional area of one VCA molecule ($A_{VCA} = \pi r_{VCA}^2$). The surface area of the GUV is $A_{GUV} = 4\pi r_{GUV}^2$. The number of VCA molecules in a GUV is given by the VCA concentration $c$ and the volume of the GUV ($N_{VCA} = c\, V_{GUV} = c\frac{4}{3}\pi r_{GUV}^3$ ). With this, we can substitute and simplify Eq. S4 to an expression that depends on the concentration of VCA in the inner buffer of the GUV:

$$S = \frac{N_{VCA}\pi r_{VCA}^2}{4\pi r_{GUV}^2}$$

$$= \frac{\pi}{3}\, r_{GUV}\, r_{VCA}^2 c \quad \text{(S5)}$$

With VCA concentrations between 0.65 and 6.5 µM, this yields surface coverages ranging from 2.5 to 25 %.

### Supporting Figures

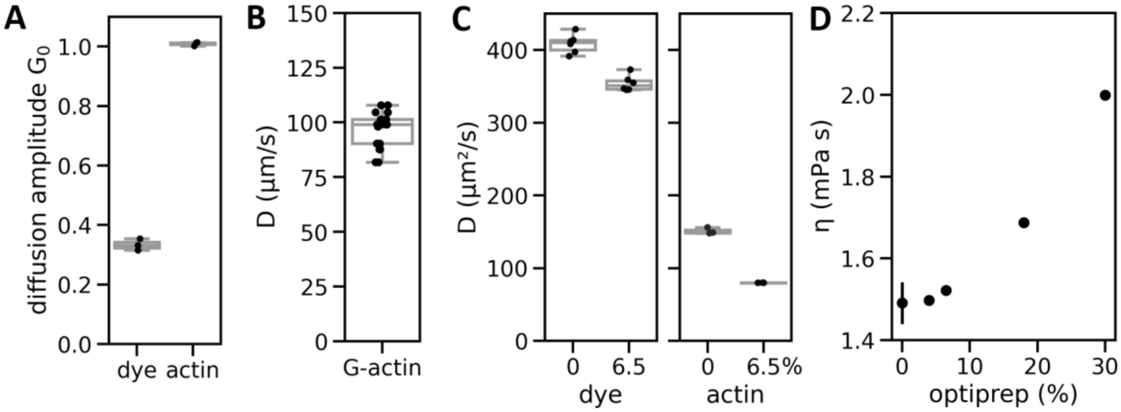


**Fig. S1. FCS on monomeric actin in buffer supplemented with Optiprep**. (A) Strip plot of the FCS diffusion amplitudes $G_0$ (see Extended Methods) for the free dye and labeled monomeric (G-) actin in a sample of AlexaFluor488-labeled G-actin in G-buffer. No extra dye was added here, so any diffusion contribution of free dye must originate from free dye contamination of the actin stock. Since the diffusion amplitude is proportional to the inverse of the concentration of the diffusing species, we conclude that there is more free dye in the sample than is bound to actin monomers, as $G_0$(dye) < $G_0$(actin). (B) Strip plot of G-actin diffusion coefficients, revealing $D$ = 96.7 ± 8.4 µm$^2$/s, N = 9 measurements from 3 separate samples. (C) Strip plots of the diffusion coefficients of free AlexaFluor488 and AlexaFluor488-labeled G-actin in G-buffer, in the presence or absence of 6.5 % Optiprep. Actin monomer diffusion is much more strongly impacted by the presence of Optiprep than the diffusion of the free dye, with the diffusion coefficient decreasing by 53 % compared to 13 % for the free dye. (D) Scatter plot of the viscosity of F-buffer supplemented with different amounts of Optiprep. Each datapoint represents the mean of 3 separate bulk rheology measurements of shear viscosity, and error bars indicate the full spread of the data. Error bars are smaller than the marker in all cases except for pure F-buffer. F-buffer viscosity increased by just 2 % upon addition of 6.5 % Optiprep.

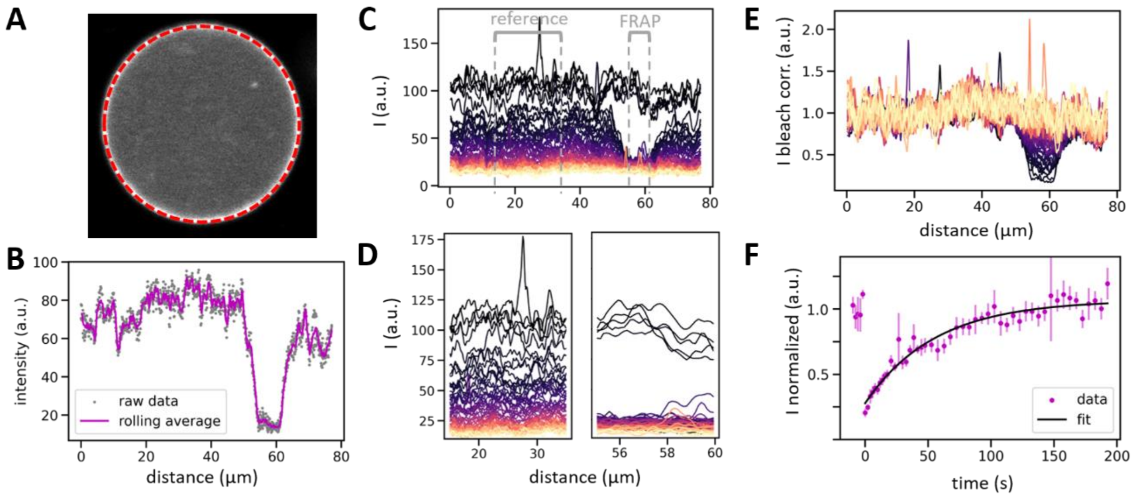


**Fig. S2. FRAP analysis procedure.** As an example, here we show a FRAP experiment assessing actin turnover in the cortex. The data analysis procedure was identical in case of FRAP measurements of the membrane or VCA diffusivity. (A) In each frame, we extract the intensity profile along the cortex (red dashed line) using the 'plot profile' function of ImageJ along a segmented line with a linewidth of 3 px. (B) The intensity profile (gray points) is smoothed by pre-processing with a rolling average over 3 data points (purple line). Here we show the intensity profile immediately after bleaching (dip around 60 µm). (C) Intensity profiles over time before correcting for bleaching. Each line represents one frame, with brighter colors indicating later times. Bleached region and reference region are indicated in gray. (D) Zoom-in on the reference (left) and FRAP ROI (right). We use the average intensity in the reference ROI to correct the whole intensity profile for bleaching in each frame. (E) Intensity profiles over time after correcting for photobleaching during image acquisition. Brighter colors indicate later times. (F) Fitting procedure. Data points show the average intensity in the FRAP region and error bars indicate the standard error of the mean intensity in the bleached region. We normalized the bleach-corrected data again such that the mean intensity before bleaching ($t$<0) was 1. The black line shows an exponential fit recovering to an asymptotic intensity $I_{inf}$. Note that fluorescence recovery due to actin monomer diffusion in the lumen was complete within the time it took to record the first frame: With a 6 µm sized FRAP region and an actin monomer diffusion coefficient of $D$ = 97 µm$^2$/s (Fig. S1B), monomer diffusion should happen with a characteristic recovery time of ~40 ms.

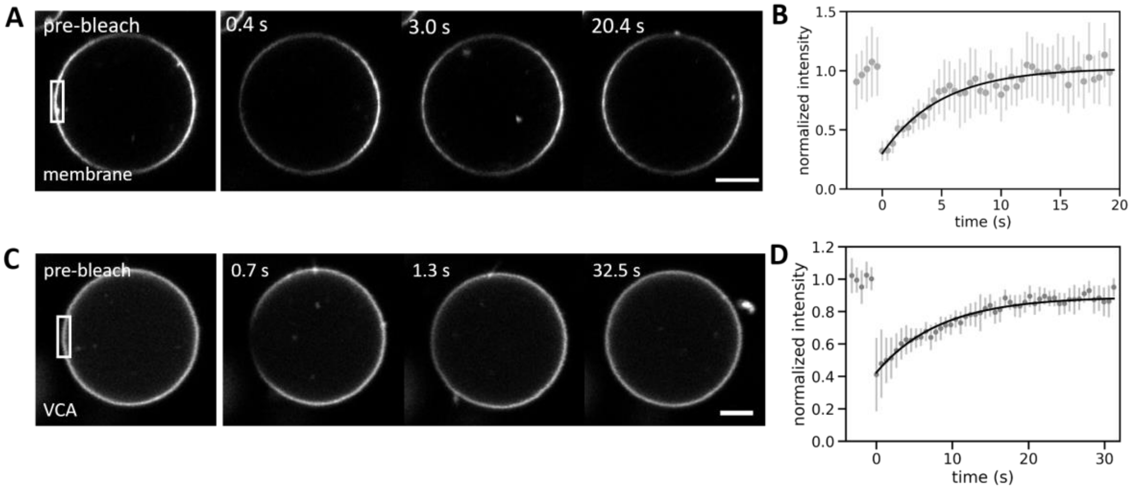


**Fig. S3. FRAP of membrane and membrane-bound VCA.** (A, C) Snapshots of a GUV membrane (A) and membrane-bound VCA (C), before and after bleaching of the ROI indicated with a rectangle in the pre-bleach images on the left. Times after photobleaching are indicated in the panels. (B, D) Symbols show recovery curves of membrane (B) and VCA (D) signal in the bleached region over time. Error bars indicate the standard deviation, black solid lines indicate exponential recovery fits with characteristic timescales of $\tau$ = 4.9 s for the membrane and $\tau$ = 8.4 s for VCA. Scale bars: 5 µm.

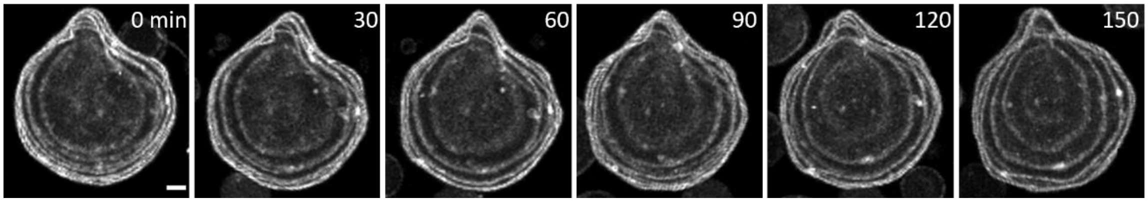


**Fig. S4. Surfaces of deflated, cortex-supported GUVs can retain their shape for several hours.** Maximum intensity projections of a GUV whose shape is stabilized by an actin cortex (shown in grey). Timelapse imaging with a framerate of two frames per hour reveals that GUV shapes can remain stable over the course of many minutes, up to several hours. To minimize photodamage to the actin cortex, the confocal z-stacks were acquired with a step height of 3 µm, spanning 33 µm in total. Scale bar: 5 µm.

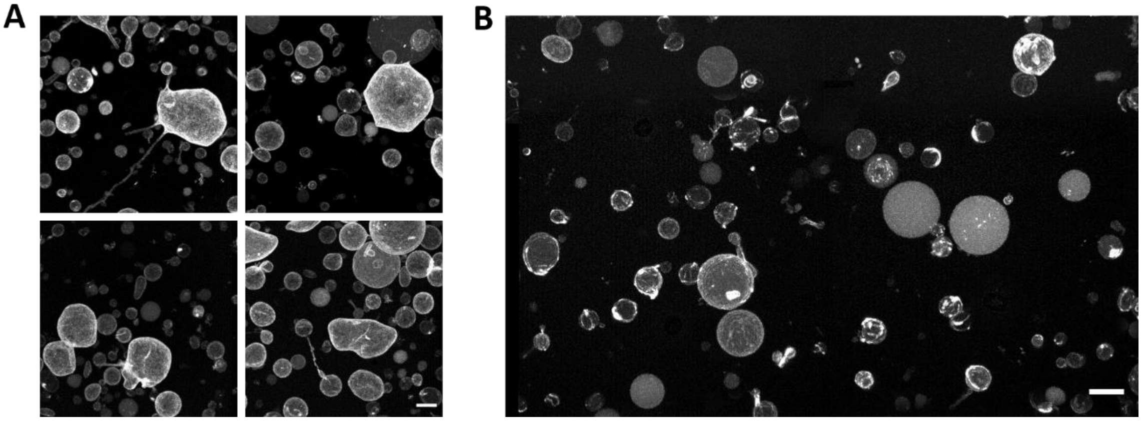

**Fig. S5. Cytochalasin D-induced actin depolymerization disassembles actin cortices and leads to spherical GUVs**. (A) Typical GUV morphologies before treatment with cytochalasin D. We observe many globally deformed GUVs and GUVs with membrane protrusions, where actin is localized strongly to the membrane. Grayscale images show actin fluorescence. (B) Typical GUV morphologies after incubating the same sample with 10 µM cytochalasin D for 90 minutes. We no longer observe globally deformed GUVs and now find actin almost exclusively in the GUV lumen. All images show maximum intensity projections of confocal stacks at 1 µm step height. Scale bars: 10 µm.

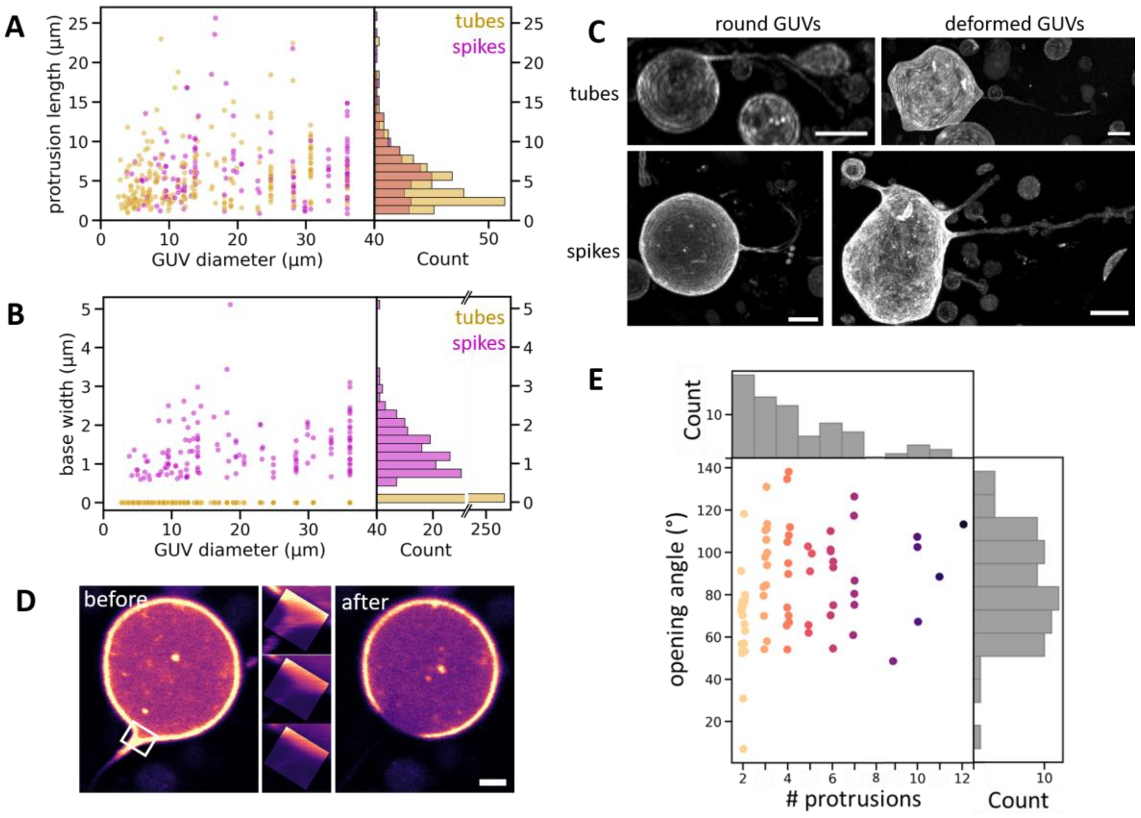


**Fig. S6. Extended characterization of actin-induced membrane protrusions of GUVs.** (A, B) Scatter plot of the length (A) and base width (B) of protrusions as a function of GUV size (left panels), together with corresponding histograms (right panels). $N$ = 261 tubes (yellow) and 157 spikes (magenta). Widths that were optically not resolvable (~ 500 nm) were set to zero. (C) Examples of GUVs with very long tubes and spikes. We found both rounded (left) and globally deformed GUVs (right) that bore protrusions longer than the GUV diameter. This was true both for tubes (top) and spikes (bottom), and we even found GUVs with multiple long protrusions (bottom right). All images show maximum intensity projections of z-stacks in the actin channel. For the top left images, we projected only the bottom half of the GUV as the long tube was otherwise obscured underneath a different GUV. All GUVs contained 8 µM actin, 6.5 µM VCA, and 50 nM ArpC1B/C5L. Scale bars: 10 µm. (D) Example of a photoablation experiment on a spike base. False-color (magma) confocal images of actin show a spike-bearing GUV, before and after photo-ablation at the base of the spike (area highlighted with white rectangle, 5.9 × 5.2 µm). Destruction of the actin cortex led to retraction of the spike (middle and right panels). Scale bar: 5 µm. (E) Scatter plot and histograms of the opening angles and number of protrusions per bouquet ($N$ = 70 bouquets on 42 GUVs). Number of protrusions is color-coded for clarity.

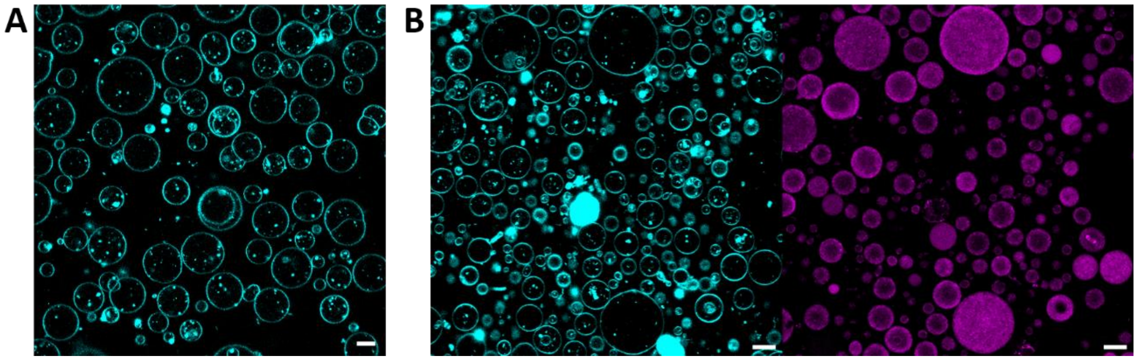


**Fig. S7. Membrane protrusions do not form in the absence of actin, or when actin polymerizes in the GUV lumen.** (A) Typical confocal image of GUVs (membrane in cyan) without encapsulated actin. (B) Typical confocal images of GUVs (membrane in cyan, left) with actin (magenta, right) spontaneously polymerizing in the lumen. The GUVs were produced in the same way as all others in this work, but VCA and Arp2/3 were omitted from the inner aqueous solution. Scale bars: 10 µm.

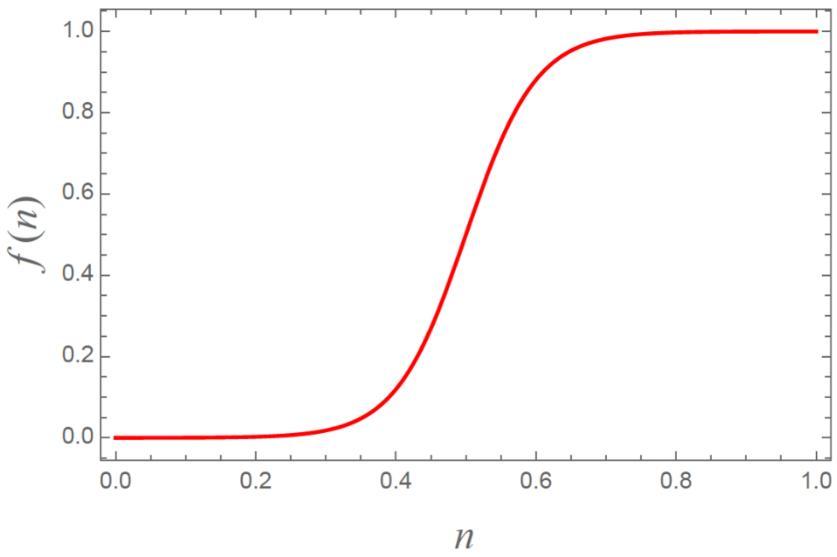


**Fig. S8. VCA response function in the mathematical model of membrane protrusion initiation.** The VCA response function *f(n)* for *a = 10* and *b = 0.5* (see Eq. S1)) reflects that some minimum local density of VCA on the surface is necessary to initiate actin polymerization, and that actin polymerization saturates above a certain VCA density.

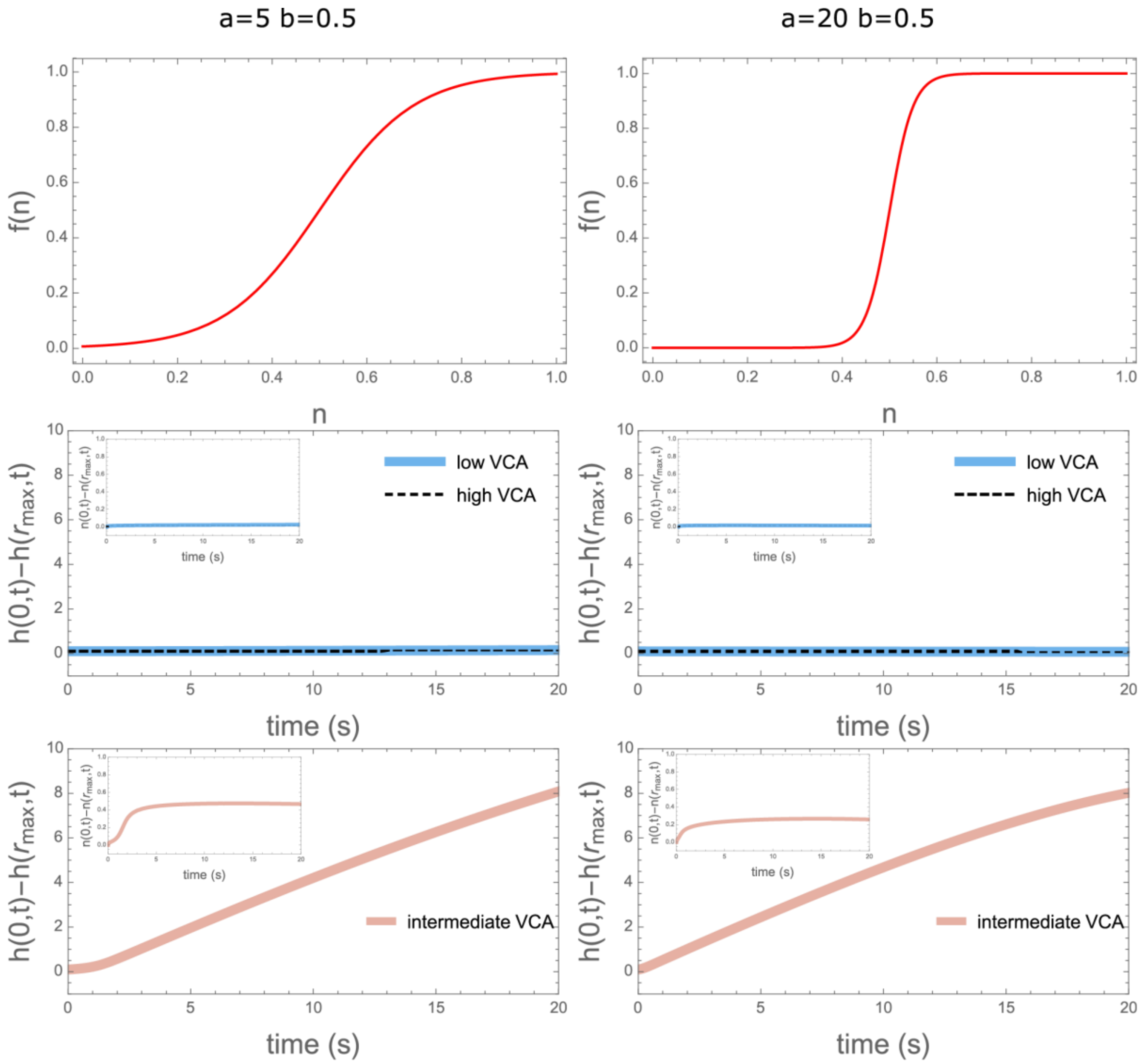


**Fig. S9. VCA response function in the mathematical model of protrusion initiation: influence of parameters *a* and *b*.** Response functions with two different sets of parameters *a* and *b* (top panels) result in comparable predictions for the membrane height evolution (middle and bottom panels) and VCA distribution (insets in middle and bottom panels).

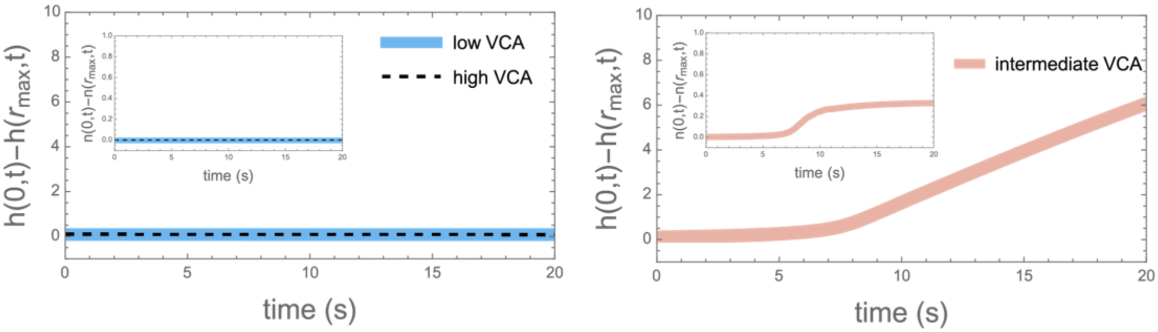


**Fig. S10. Computational results are robust against changes to the parameter α.** The mathematical model still predicts no protrusion initiation for high and low VCA density (left) and protrusion initiation for intermediate VCA density (right) even when α is reduced 10-fold from 0.1 to 0.01.

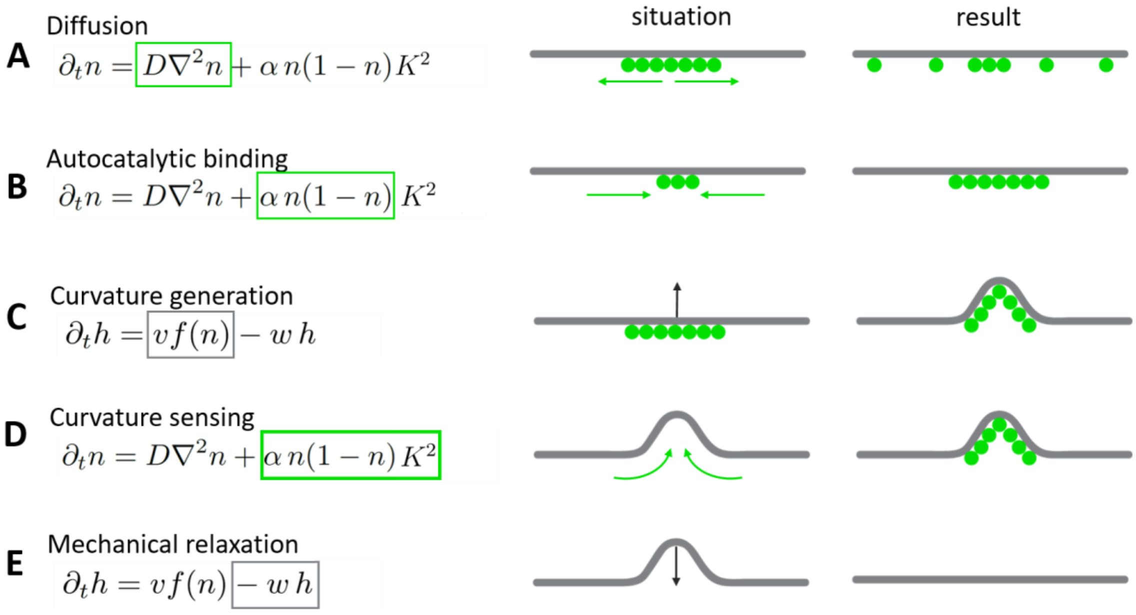


**Fig. S11. Interplay between membrane deformation and VCA density in the model of protrusion formation.** Schematic representations of the interaction terms in the model, illustrating how the different terms drive the evolution of the membrane position (gray in the images on the right) and VCA surface density (green in the images). (A) Diffusion smears out concentrated regions of VCA along the membrane. (B) Autocatalytic binding, modeled after the autocatalytic growth of Arp2/3 nucleated actin networks, means that new VCA accumulates preferentially where VCA is already bound. (C) Curvature generation depends on VCA density: Where there is a lot of VCA (and thus actin polymerization), membranes are displaced and thus bent. (D) Curvature sensing emerges because a membrane deformation implies the presence of actin and thus mother filaments, with which VCA can interact to generate new daughter filaments. Via actin, VCA is thus recruited to locations where the membrane is curved. (E) Mechanical relaxation occurs since a deformed membrane experiences a restoring force driving it back to a flat conformation.

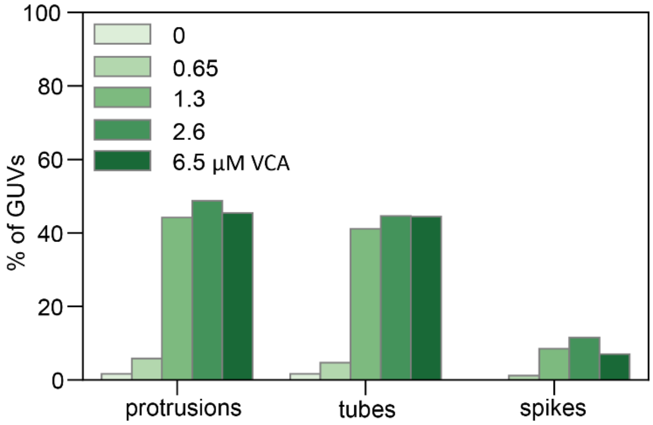


**Fig. S12. Spikes are more strongly suppressed at high VCA concentrations than tubes.** At high VCA surface densities, the prevalence of tubes stays virtually unchanged but spikes are significantly reduced.

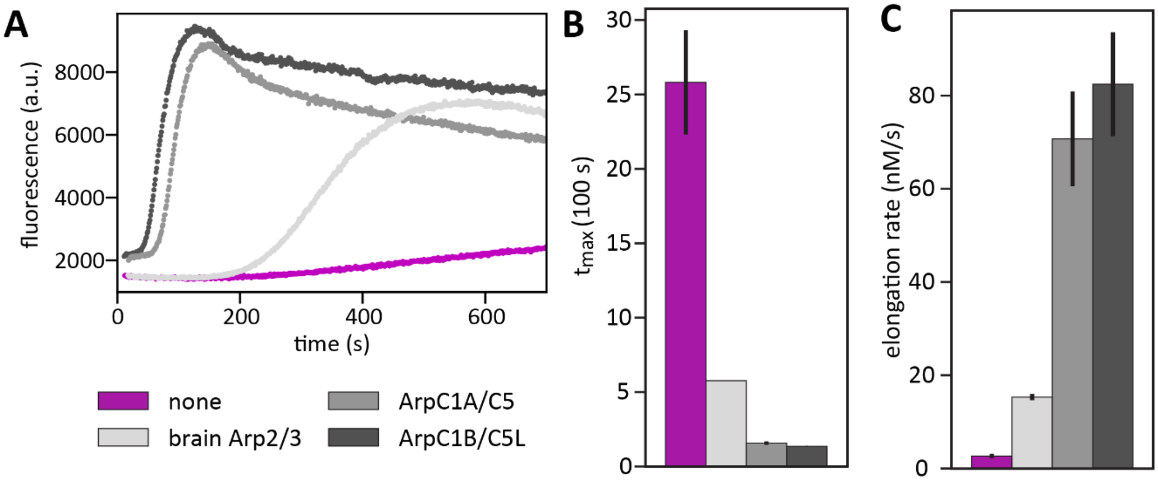


**Fig. S13. Pyrene fluorescence-based quantification of actin polymerization by different human Arp2/3 isoforms and by Arp2/3 from different species.** (A) Typical examples of pyrene polymerization curves of actin without nucleating proteins (magenta) or with different isoforms of Arp2/3 (shades of gray). (B, C) Time after which the maximum pyrene fluorescence was reached, and elongation rate at the time when half of all actin was polymerized. Bars and error bars denote the average and SEM of 2 independent measurements for each Arp2/3 isoform, and 4 independent measurements for actin alone. Quantitative analysis of the polymerization curves reveals that, while all Arp2/3 isoforms speed up actin polymerization significantly compared to spontaneous actin polymerization, commercially available Arp2/3 isolated from porcine brain was around a factor of 5 less effective at promoting actin polymerization than the recombinant human isoforms ArpC1A/C5 and ArpC1B/C5L produced in insect cells**.**

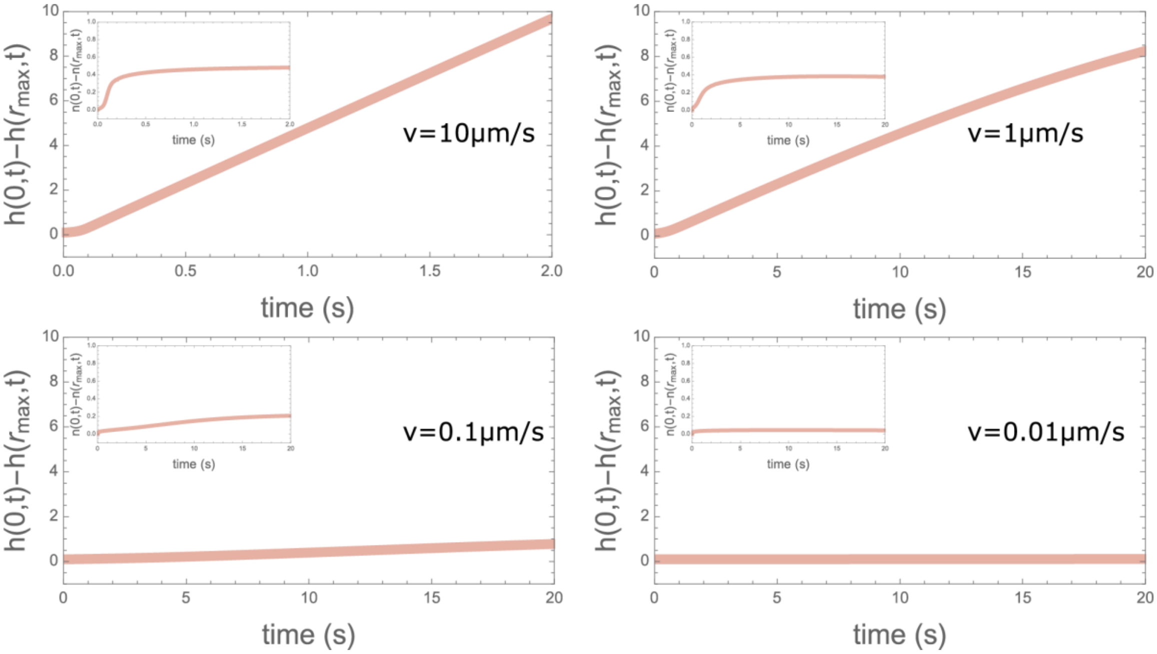


**Fig. S14. Mathematical modeling predicts that membrane protrusions form consistently over a wide range of actin polymerization velocities.** Main plots: Relative membrane height at the origin as a function of time for a constant, intermediate concentration of VCA and for different actin polymerization velocities ranging from 0.01 to 10 µm/s (see legends). Insets: Relative VCA distribution at the origin as a function of time. Protrusions grow more rapidly when actin polymerization is fast, but protrusions are still initiated for polymerization velocities as low as 0.1 µm/s.

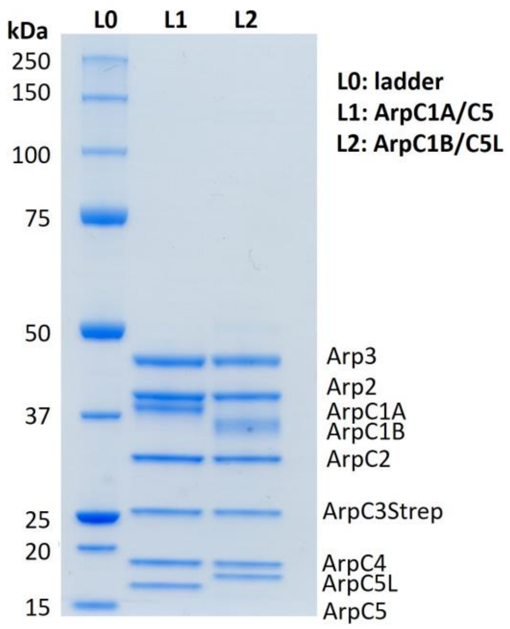


**Fig. S15.** SDS PAGE gel of purified human Arp2/3 isoforms. Lane 0 contains a protein ladder for reference, with protein molecular weights (in kDa) shown on the left. The two human Arp2/3 isoforms, with ArpC1A/C5 in lane 1 (L1) and ArpC1B/C5L in lane 2 (L2), are pure and all the subunits are stoichiometric (see legend on the right).

**Table S1. Summary of parameters in the mathematical model of actin-mediated membrane protrusion initiation.**

| Parameter | Value used | Reference |
|---|---|---|
| VCA diffusion constant $D$ | 1 μm$^2$/s | (14) |
| Actin maximum polymerization velocity $v$ | 1 μm/s | (14) |
| Rate of relaxation of membrane deformations $w$ | 0.013 s$^{-1}$ | (1) |
| VCA enrichment constant on the membrane $\alpha$ | 0.1 μm$^2$/s | - |
| VCA response function parameter $a$ in Eq. S1 | 10 | - |
| VCA response function parameter $b$ in Eq. S1 | 0.5 | - |

**Table S2. Confocal imaging settings.** The left column lists compounds and their fluorescent labels, either AlexaFluor488 or Cy5. The Excitation column lists excitation wavelength, laser attenuation, and exposure time (camera-based microscope) or pixel dwell time (point scanning microscope).

| Compound | Instrument | Excitation | Detection |
|---|---|---|---|
| actin-AlexaFluor488 | Olympus spinning disk | 491 nm laser, 10.5 %, 200 ms exposure time | Andor iXon X3 EM-CCD, gain 250 |
| membrane-Cy5 | Olympus spinning disk | 640 nm laser, 53 %, 200 ms exposure time | Andor iXon X3 EM-CCD, gain 250 |
| actin-AlexaFluor488 | Leica Stellaris LSCM | WLL at 499 nm, 18 %, 3.16 μs pixel dwell time | HyDS2 (504-590 nm), counting mode |
| membrane-Cy5 | Leica Stellaris LSCM | WLL at 640 nm, 2 %, 3.16 μs pixel dwell time | HyDX3 (658-809 nm), counting mode |
| VCA-AlexaFluor488 | Leica Stellaris LSCM | WLL at 499 nm, 0.5 %, 3.16 μs pixel dwell time | HyDX1 (510-567 nm), Standard mode, gain 60 |

**Table S3. Laser settings for photobleaching and –ablation experiments.** Laser settings listed here refer to the settings used to bleach fluorophores or ablate actin structures. Imaging settings for the pre- and post-bleach images are listed in table S1.

| **Compound** | **Laser settings** | **Timing** |
|---|---|---|
| actin-AlexaFluor488 (FRAP) | WLL at 483, 491 and 499 nm, 100 % each, 3.16 µs pixel dwell time | Pre-bleach: 5 frames at 0.51 fps. Post-bleach: 10 frames at 0.51 fps, then 10 frames at 0.33 fps, and finally 30 frames at 0.2 fps |
| membrane-Cy5 (FRAP) | WLL at 643, 648, 653 and 658 nm, 100 % each, 0.95 µs pixel dwell time | Pre-bleach: 5 frames at 4 fps. Post-bleach: 50 frames at 4 fps |
| VCA-AlexaFluor488 (FRAP) | WLL at 470, 483, 491 and 499 nm, 100 % each, 6.34 µs pixel dwell time | Pre-bleach: 5 frames at 1.54 fps. Post-bleach: 50 frames at 1.54 fps |
| actin (photoablation) | 1.6 W 405 nm solid state laser, 100 %, 6.24 µs pixel dwell time. 4 rounds for local ablation, 8 rounds for ablation of the entire GUV cortex. | Single z-stack (step size 1 µm) before and after ablation. |

**Movie S1 (separate file).** Movie of the equatorial confocal slice of a GUV with no actin cortex (left panel in Fig. 2 B), showing its deformations over the course of 50 s. Scale bar: 5 μm.

**Movie S2 (separate file).** Movie of the equatorial confocal slice of a cortex-bearing GUV (right panel in Fig. 2 B), showing its deformations over the course of 50 s. Scale bar: 5 μm.

**Movie S3 (separate file).** More examples of cortex-supported GUVs over 50 s, showing that actin cortices suppressed shape fluctuations. Scale bar: 10 μm.

**SI References**

1. N. S. Gov, A. Gopinathan, Dynamics of membranes driven by actin polymerization. *Biophys. J.* **90**, 454–469 (2006).
2. C. Simon, *et al.*, Actin dynamics drive cell-like membrane deformation. *Nat. Phys.* **15**, 602–609 (2019).
3. J. Weichsel, P. L. Geissler, The more the tubular: Dynamic bundling of actin filaments for membrane tube formation. *PLoS Comput. Biol.* **12**, e1004982 (2016).
4. J. Neuhold, *et al.*, GoldenBac: A simple, highly efficient, and widely applicable system for construction of multi-gene expression vectors for use with the baculovirus expression vector system. *BMC Biotechnol.* **20**, 26 (2020).
5. E. P. Petrov, T. Ohrt, R. G. Winkler, P. Schwille, Diffusion and segmental dynamics of double-stranded DNA. *Phys. Rev. Lett.* **97**, 258101 (2006).
6. J. Widengren, R. Rigler, Fluorescence correlation spectroscopy of triplet states in solution: a theoretical and experimental study. *J. Phys. Chem.* **99**, 13368–13379 (1995).
7. L. Baldauf, F. Frey, M. A. Perez, T. Idema, G. H. Koenderink, Reconstituted branched actin networks sense and generate micron-scale membrane curvature. *biorxiv Prepr.*, 2022.08.31.505969 (2022).
8. L. K. Doolittle, M. K. Rosen, S. B. Padrick, "Measurement and analysis of in vitro actin polymerization" in *Methods in Molecular Biology*, A. S. Coutts, Ed. (Springer Science and Business Media LLC, 2013), pp. 273–293.
9. H. A. Faizi, C. J. Reeves, V. N. Georgiev, P. M. Vlahovska, R. Dimova, Fluctuation spectroscopy of giant unilamellar vesicles using confocal and phase contrast microscopy. *Soft Matter* **16**, 8996–9001 (2020).
10. M. D. Flanagan, S. Lin, Cytochalasins block actin filament elongation by binding to high affinity sites associated with F-actin. *J. Biol. Chem.* **255**, 835–838 (1980).
11. P. Forscher, S. J. Smith, Actions of cytochalasins on the organization of actin filaments and microtubules in a neuronal growth cone. *J. Cell Biol.* **107**, 1505–1516 (1988).
12. M. Fritzsche, A. Lewalle, T. Duke, K. Kruse, G. Charras, Analysis of turnover dynamics of the submembranous actin cortex. *Mol. Biol. Cell* **24**, 757–767 (2013).
13. H. Fischer, I. Polikarpov, A. F. Craievich, Average protein density is a molecular-weight-dependent function. *Protein Sci.* **13**, 2825–2828 (2004).
14. N. S. Gov, Dynamics and morphology of microvilli driven by actin polymerization. *Phys. Rev. Lett.* **97**, 018101 (2006).
